# Rapid evolution of *Klebsiella pneumoniae* biofilms *in vitro* delineates adaptive changes selected during infection

**DOI:** 10.1101/2024.03.16.585345

**Authors:** Greta Zaborskytė, Patrícia Coelho, Marie Wrande, Linus Sandegren

## Abstract

Biofilm formation facilitates infection by the opportunistic pathogen *Klebsiella pneumoniae*, primarily through indwelling medical devices. Here, we address how *K. pneumoniae* can increase its biofilm capacity by experimental evolution of surface-attached biofilms to mimic catheter-associated infections. We observed rapid convergent evolution that altered or abolished capsule, modified the fimbrial adhesin MrkD, or increased production of fimbriae and cellulose via upregulated c-di-GMP-dependent pathways. However, evolutionary trajectories and resulting phenotypes showed strain differences, illustrating the importance of genetic background on biofilm adaptation. Multiple biofilm aspects, such as early attachment, biofilm topology, surface preference, and extracellular matrix composition, were affected in a mutation-specific manner. Acute virulence was linked to the underlying genetic change rather than the overall biofilm capacity. Single mutations conferring hypermucoidy and c-di-GMP-related changes extensively overlapped with previously identified adaptive changes in UTI and wound isolates, confirming biofilm as an important selective trait *in vivo*.

## Introduction

Biofilm growth is an inherent component of opportunistic infections^1^. Indwelling medical devices that support biofilm growth often facilitate the transition between colonization and infection^2,3^. For classical *K. pneumoniae,* this translates into hospital-acquired urinary tract infections (UTI) or ventilator-associated pneumonia exacerbated by the presence of urinary catheters or endotracheal tubes^4–7^. Nevertheless, the success of opportunistic *K. pneumoniae* as a nosocomial pathogen is often studied from the perspective of its multidrug resistance^8^, whereas its biofilm growth that enables both patient colonization and survival in hospital settings remains understudied.

During colonization or infection, bacterial populations experience selection to adapt to different host niches^9^. Increased biofilm capacity is often found as a within-host adaptation throughout prolonged chronic infections, for example, for *Pseudomonas aeruginosa* and other bacteria in cystic fibrosis lungs^10,11^. Since biofilm communities embedded in extracellular matrix are much more recalcitrant to antibiotic treatment and immune factors^1^, they do not experience the same host-related bottlenecks as non-biofilm populations. Therefore, such populations can be maintained for extended periods of time and serve as a reservoir of bacteria. Furthermore, due to spatial constraints, biofilm growth results in formation of distinct microenvironments that support further diversification^12,13^. This is reflected in long-term within-host adaptations during biofilm-associated infections, where populations diversify on many other aspects important for survival inside the host, such as iron scavenging or antibiotic resistance^10,14,15^. Therefore, from an eco-evolutionary perspective, changes in biofilm capacity play an essential role in the overall pathoadaptive landscape of infection sites. To elucidate the molecular mechanisms behind such adaptations, experimental evolution in different biofilm model systems has proven a powerful tool^16,17^.

While *K. pneumoniae* can asymptomatically colonize the gut for extended periods of time, the infections caused by this bacterium are rarely chronic. This is probably due to its medical-device related infection nature, where the indwelling medical devices are removed upon infection, and therefore, population maintenance is different from biofilms that preferentially form on biological surfaces. However, during our recent study on extensive clonal *K. pneumoniae* outbreak, we observed a repeated mutation-mediated switch to a chronic, rather than acutely virulent, state in the infection sites that often involved catheterization^18^. We also identified increased biofilm formation on catheter-like surfaces as an adaptive phenotype repeatedly selected in patients mostly with urinary tract infections^18^. Therefore, the question arises whether such adaptations are selected due to the presence of an abiotic surface or are co-selected with other phenotypes.

Here, we have explored evolutionary trajectories and the underlying genetic networks leading to increased biofilm formation in three clinical *K. pneumoniae* strains, including the clone from the hospital outbreak. Mutants with much better biofilm capacity were rapidly selected during short-term experimental evolution in an *in vitro* biofilm system mimicking catheter surfaces. Genotypically, there was an extensive convergence among the strains and independent lineages. Single nonsynonymous mutations were enough to drastically change the phenotypes in diverse but highly specific mutation- and strain-dependent ways. We also show that the biofilm-increasing mutations cause possible trade-offs in other features relevant during infection, such as sensitivity to innate immune factors. Importantly, the *in vitro* selected changes strongly resemble those selected during within-host evolution and provide insights into why such changes can be advantageous during infection.

## Results

### Increased biofilm capacity is rapidly selected on catheter-like surfaces via mutations and morphotypes that match those selected clinically

To explore how *K. pneumoniae* evolves increased biofilm formation on abiotic surfaces, we performed experimental evolution in the FlexiPeg biofilm model^19^ (Fig. 1a). We initially evolved the bacteria on uncoated surfaces, but to make the conditions resemble the clinical situation during catheterization, we coated the pegs with silicone and further with fibrinogen from human plasma. Fibrinogen has been shown to heavily coat inserted catheters as a result of the local inflammatory response^20^ and thus promote biofilm formation in some pathogens^21,22^ including *K. pneumoniae*^6^. To evaluate the effect of genetic background on evolutionary trajectories, we used three clinical *K. pneumoniae* strains of the classical (opportunistic) pathotype: the index isolate DA14734 from a large clonal hospital outbreak in Uppsala, Sweden^23,24^, the UTI isolate C3091^25^, and the respiratory isolate IA565^26^. The isolates belong to different sequence-, capsule-, and O-antigen types (*see Materials and Methods*).

**Fig. 1.**
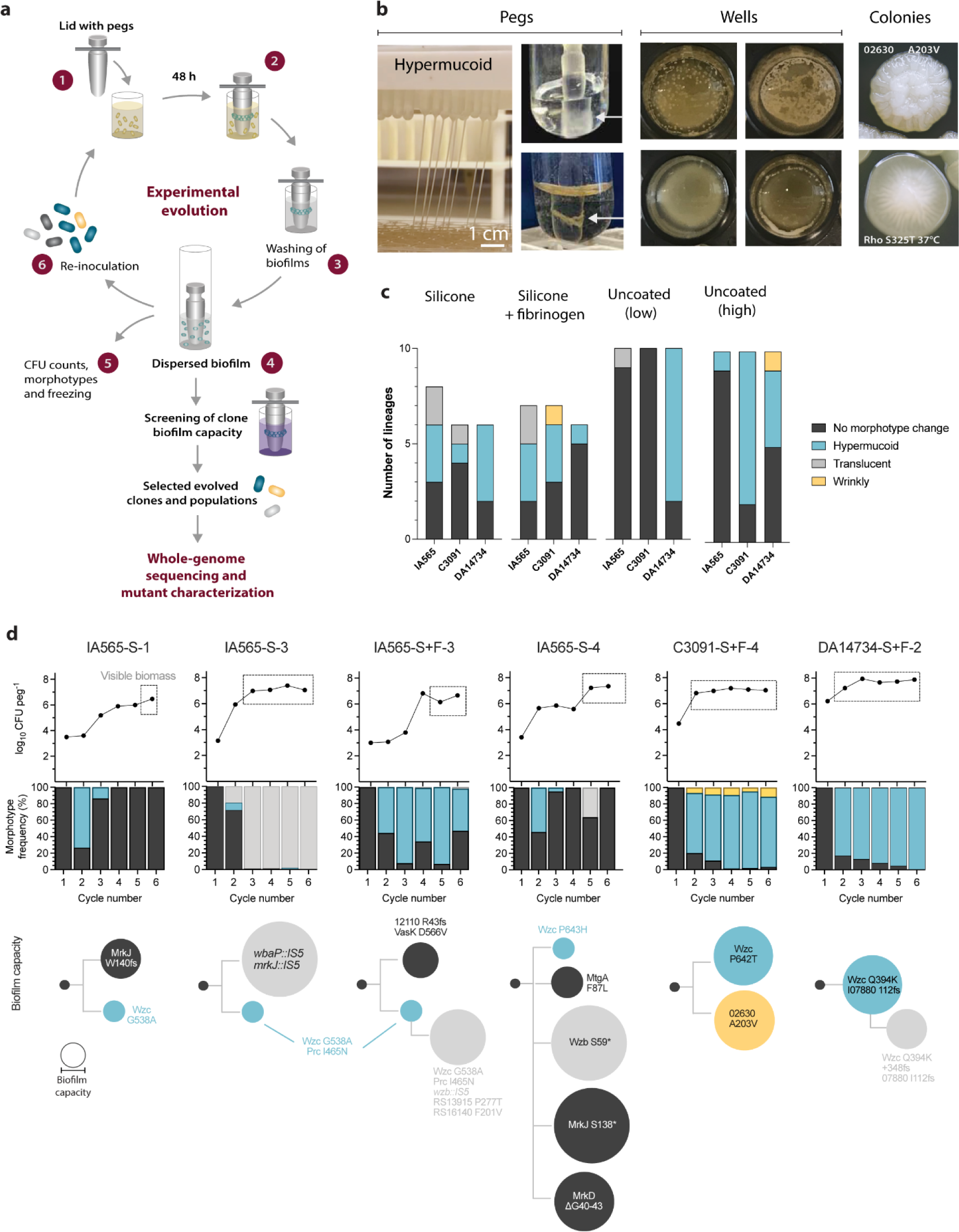
Fast convergent evolution of increased biofilm formation in different *K. pneumoniae* strains. **a,** Scheme of experimental evolution of *K. pneumoniae* IA565, C3091 and DA14734 on the FlexiPeg device. (1} Diluted cultures were added to a 96-well plate and allowed to form biofilms on pegs for 48 h (2) with a medium change after 24 h. (3) Pegs with attached biofilms were washed to remove planktonic and loosely bound bacteria, and (4) biofilms were dispersed by vortexing. (5) CFU/peg and morphotype frequencies in each lineage were assessed after each cycle. (6) Part of the harvested biofilm population was used to re-inoculate new biofilms. **b,** Examples of biofilm phenotypes on pegs, in the wells, and the two wrinkly colony morphologies, **c,** Quantification of evolving lineages with respect to morphotype (colony morphology) changes during cycling. Uncoated low and high refer to the size of the population transferred (0.25% and 25%, respectively) during cycling on pegs without silicone coating, **d,** Examples of evolutionary trajectories. Biofilm population size (top dot plots), frequency of different morphotypes (middle bar graphs), evolutionary trajectories, and biofilm capacity (CV-stained biomass of 48 h biofilms) of whole-genome sequenced clones (bottom panels).

Evolving biofilm populations displayed gradual or stepwise increases in biofilm population size of up to 4-log CFU/peg, mostly coinciding with a substantial accumulation of biomass on the pegs, lineage-specific aggregations in the wells, and appearance of specific colony morphologies (morphotypes) (Fig. 1 b-d, Supplementary Fig. 1 and Data 1, and Supplementary Videos 1-5). Biofilm formation was generally more robust on uncoated pegs and in the presence of fibrinogen, and the maintenance of biofilm populations was more stable on these surfaces, especially for the wound isolate DA14734 (Supplementary Fig. 1). However, mutants selected on a smooth silicone surface on average evolved a higher biofilm capacity than those in lineages evolving with fibrinogen (Fig. 2a), suggesting higher evolvability under less optimal conditions. While single nonsynonymous point mutations were enough to increase biofilm capacity, combinations of mutations were also selected, and they were more common in clones on silicone or silicone with fibrinogen than uncoated (Fig. 2 b and c). These clones also had a more diverse mutation spectrum than those selected on uncoated surfaces, especially for the respiratory IA565 and UTI C3091 strains (Fig. 2d). There was no apparent bias of specific mutations for any condition. No trade-off in planktonic growth rate was seen for the evolved mutants (Supplementary Fig. 2).

**Fig. 2.**
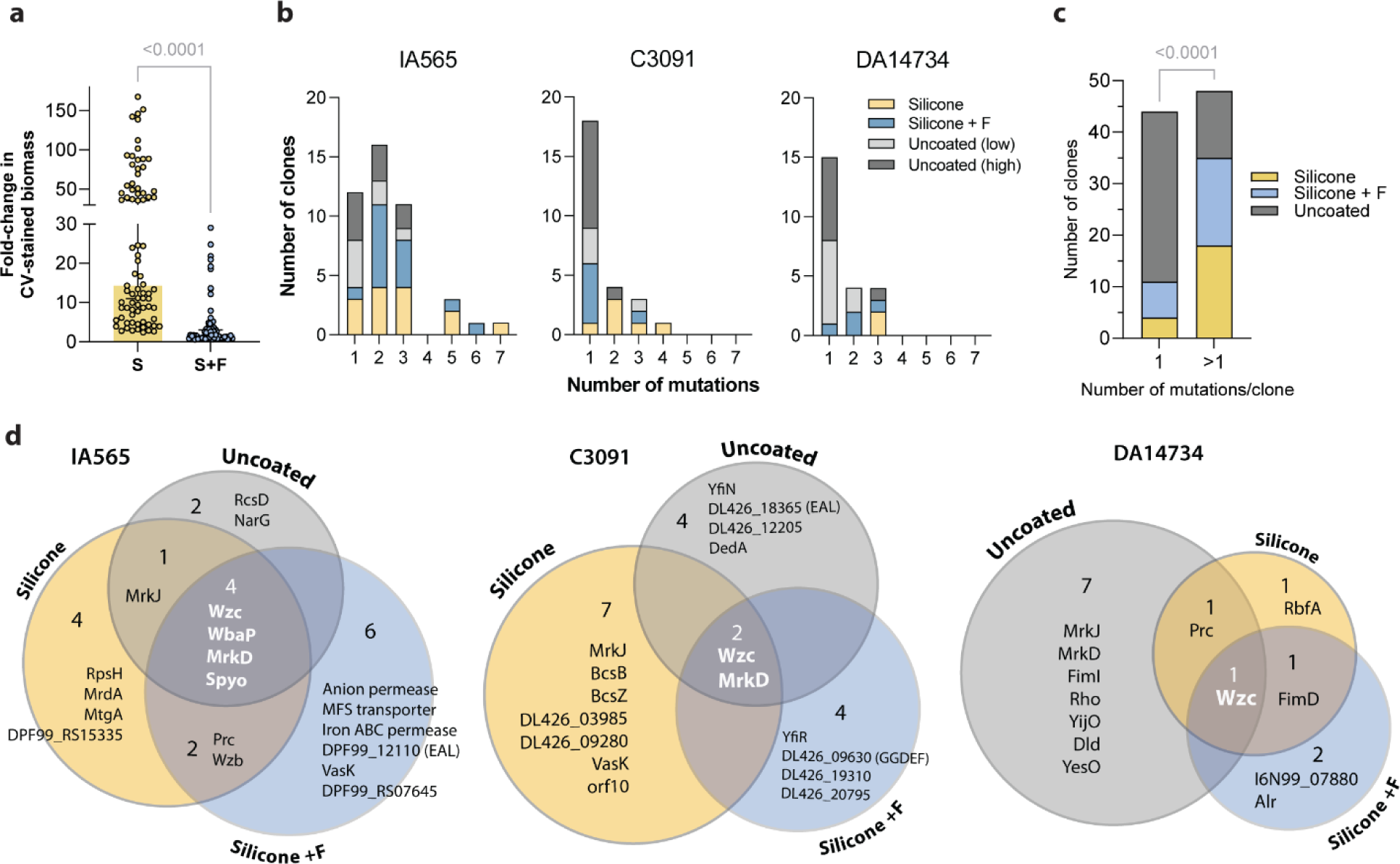
The influence of biofìlm growth surface on mutational characteristics. **a,** Biofilm capacity of clones evolved on silicone or silicone with fibrinogen. Each datapoint shows a single clone included in the screening after the cycling. The biofilm capacity of each clone was measured in the same conditions as during cycling, S - silicone, S+F - silicone with fibrinogen. The values are relative to the respective parental strain. The comparison between different surfaces was done by the Mann-Whitney test, **b,** Histograms showing the number of mutations in individual evolved clones, **c,** Distribution of clones with single mutations or mutation combinations in lineages evolved on different surfaces. Statistical significance tested by Chi-square, **d,** Venn diagrams showing the overlap of mutated genes after selection on different surfaces for IA565, C3091, and DA14734 strains.

The most common morphotypes linked to increased biofilm formation were hypermucoid or translucent colonies, indicating altered capsule production^27,28^. The hypermucoid morphotype was mediated by missense mutations in *wzc*, encoding a conserved tyrosine autokinase that is an essential part of the Wzy-dependent capsule synthesis and export machinery in Gram-negative bacteria^29–31^. Capsule-loss mutants displayed a translucent morphotype due to inactivation of the initial glycosyltransferase WbaP or the phosphatase Wzb, or were derived from the hypermucoid morphotype by additional frameshifts in *wzc*. In two lineages, a wrinkly/rugose morphotype occurred (Fig. 1b). The genetic basis for one of these variants was a point mutation in an uncharacterized GGDEF domain protein gene (DL426_09630^A203V^), possibly involved in c-di-GMP biosynthesis or sensing. The second wrinkly phenotype was temperature dependent (wrinkly at 37℃, parental-like at 30℃) and mediated by an S325T mutation in the global transcription terminator Rho. Different morphotypes could sweep, replace each other, or co-exist over time during evolution (Fig. 1d and Supplementary Data 1). The Wzc-mediated hypermucoid morphotype frequently dominated the biofilm population early during cycling for C3091 and DA14734. In contrast, this morphotype was often replaced by others for IA565, such as a translucent (non-capsulated) morphotype. However, this clonal interference depended on the composition of the evolving population and possibly the surface. One lineage showed an interesting maintenance of both hypermucoid and wrinkly morphotypes at the same frequencies throughout all six cycles, suggesting a possible cooperative interaction (e.g., due to niche construction). However, change in morphotype was not a prerequisite for increased biofilm capacity since more than half of the lineages (56/96) had no morphotype changes despite accumulating biofilm-increasing mutations (Fig. 1c and Supplementary Table 1).

Capsule, type 3 fimbriae, and c-di-GMP-dependent signaling were repeatedly targeted in independent lineages and across the three parental strains, especially the Wzc tyrosine autokinase (33/96 lineages), the MrkD fimbrial adhesin (20/96 lineages), and the MrkJ phosphodiesterase (12/96 lineages) (Fig. 3 and Supplementary Table 1). An uncharacterized gene in the IA565 strain repeatedly acquired the same mutation, T361S, and is followed up in another study. Notably, there was a substantial overlap in functional targets, individual genes, and even mutated amino acid positions between biofilm-mutants selected *in vitro* and isolates from the clonal hospital outbreak^18^ (Fig. 3). The overlapping targets/mutations were primarily found in outbreak isolates from infection sites, especially UTIs, and not as often in colonizing isolates (two-tailed binomial test, p<0.0001). The hypermucoid *wzc* mutants and I6N99_05140 (DL426_09630)^A203V^ mutation conferring a wrinkly phenotype were selected exclusively during infections (UTIs and wound), suggesting that they represent niche-specific adaptations where biofilm formation may be advantageous. Together, these results illustrate that *K. pneumoniae* can rapidly diversify when grown on surfaces to increase biofilm capacity and that the selected genotypes are highly convergent across strains and resemble adaptations selected during human infections.

**Fig. 3.**
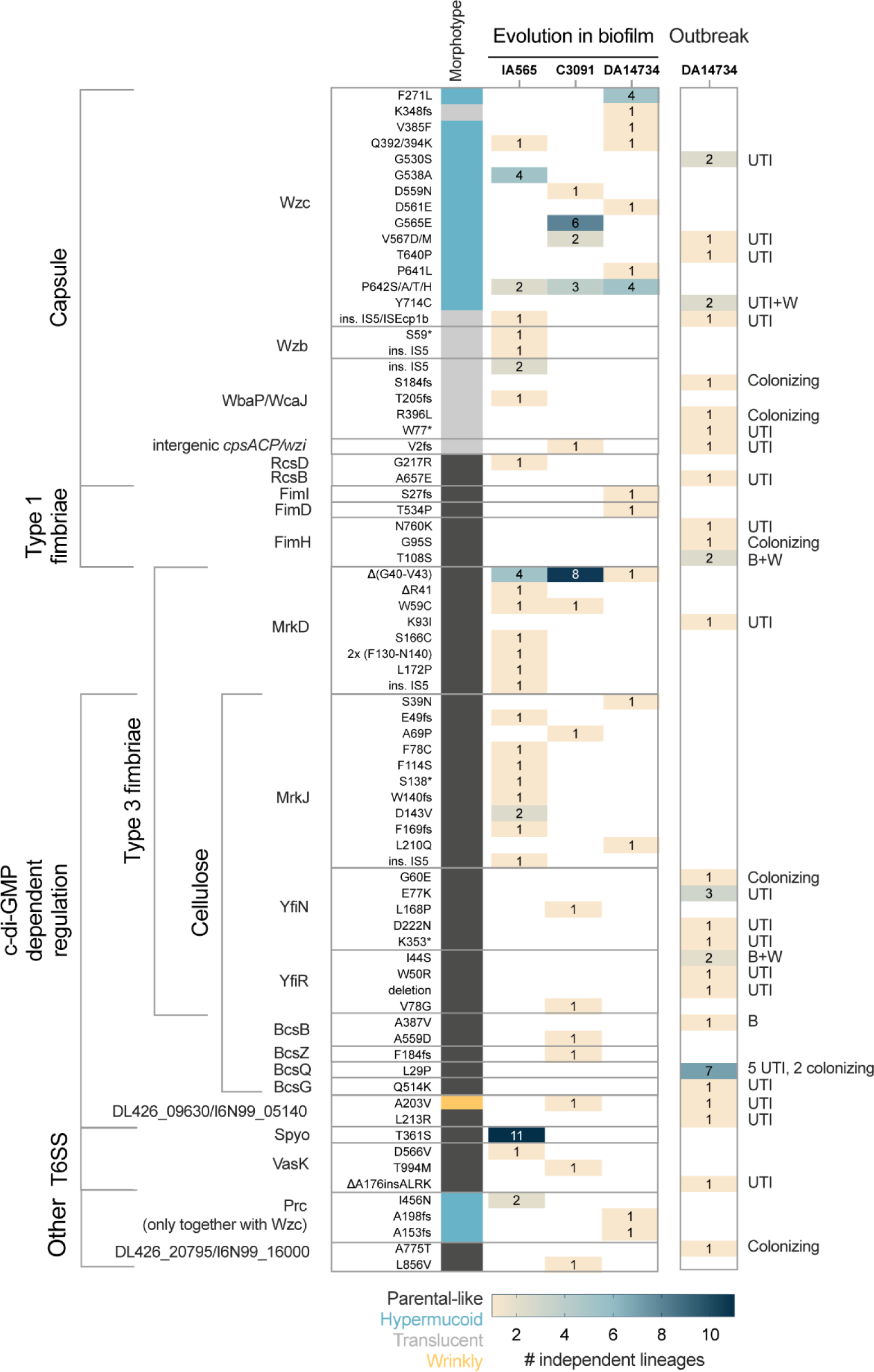
Functional grouping of mutations repeatedly selected alone or in combination with other changes. The numbers in columns IA565, C3091, and DA14734 indicate how many independent lineages from all cycling rounds combined (uncoated, silicone, silicone+fibrinogen) had clones with the respective mutation. Column “Outbreak” refers to how many independent isolates had the respective mutation during the outbreak at Uppsala University Hospital^13^. Isolates from the outbreak are marked as colonizing (faecal screening) and from infections: urinary tract (UTI), blood (B), and wound (W).

### Various levels of mucoviscosity are advantageous in biofilm but confer different phenotypes

Capsule was the most common target during biofilm evolution in all strains, although the frequency of the types of mutations selected (Fig. 3) and their evolutionary dynamics in populations differed (Fig. 1d). The Wzc tyrosine autokinase was the most frequently targeted gene among evolved lineages. The selected point mutations overlapped among all strain backgrounds and targeted conserved residues in the periplasmic and cytoplasmic regions, including the kinase domain (Fig. 4a). However, the phenotypic properties conferred by these mutations depended on the strain background, which is likely attributed to the different capsule types of the strains: *wzc* mutant colonies in IA565 (KN2) produced classical “strings” in the string test, while those of C3091 (K16) and DA14734 (K51) were completely stuck on the agar (Supplementary Videos 2-5). Wzc mutants in IA565 were not as hypermucoid in liquid as those of C3091 and DA14734 (Fig. 4b). The biofilm phenotype of *wzc* missense mutants in C3091 and DA14734 on the pegs was exceptional with biomass that stretched >10 cm from the pegs after lifting the peg lid (Fig. 1b and Supplementary Video S1), while *wzc* mutants in IA565 formed less biomass on the pegs (Fig. 4c). Accordingly, SEM images of *wzc* mutants in C3091 clearly showed more biofilm than in the IA565 background (Fig. 4d). However, regardless of the strain background, *wzc* mutants did not attach to polystyrene (Fig. 4c) but formed a lot of “fluffy” non-surface attached biomass in the wells, which might explain why a previous study reported *wzc* mutants as non-biofilm forming^28^.

**Fig, 4.**
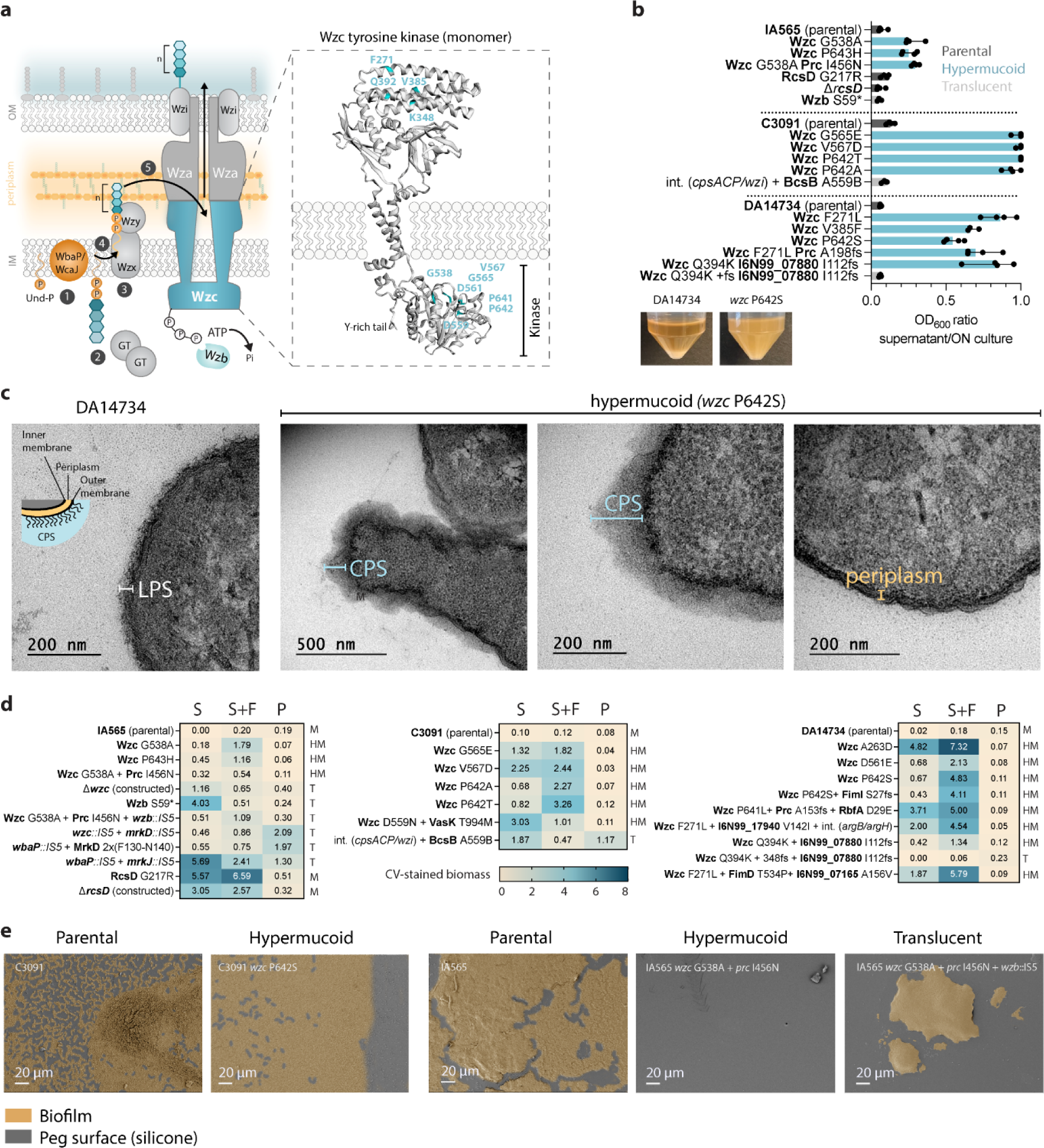
Characteristics of capsule mutants. **a.** Schematic representation of capsule synthesis in *K, pneumoniae* focusing on the Wzc protein and mutations selected during experimental evolution. The Wzc structure is a monomer (chain A) from *E. coli* (PDB 7NHR). **b** Sedimentation assay showing the ratio between optical density (600 nm) of the supernatant after low-speed centrifugation versus before centrifugation. Dots represent four biological replicates. Blue bars show hypermucoid *wzc* mutants, dark grey - parental-like colony morphology, light grey - non-mucoid (translucent). Images below show parental DA14734 (left) and wzcP642S (right) cultures after low-speed centrifugation at 1000xg for 5 min. **c,** CV-stained biofilm biomass on silicone pegs (S), silicone + fibrinogen pegs (S+F), and in polystyrene wells (P) after 48 h. Heat map values illustrate medians of at least four biological replicates. Morphotype is indicated for each mutant as mucoid (M), hypermucoid (HM), and translucent (T). **d,**Transmission electron micrographs of the DA14734 parental strain and the *wzc*^P642S^ mutant, **e,** SEM images of 48 h biofilms formed by capsule mutants on silicone-coated pegs. Raw images have been colour-modified to distinguish biofilm-covered areas.

We looked further into what effects the *wzc* mutations had on a cellular level. A constructed in-frame deletion of *wzc* led to non-mucoid colonies, confirming that the selected *wzc* missense mutations were not inactivating but altered the function of Wzc. Polysaccharide extracts from *wzc* mutants were extremely viscous after ethanol precipitation, in contrast to the compact pellet from the parental strains, indicating more and possibly longer polysaccharide molecules in the mutants, as shown for *wzc* missense mutants in *A. baumannii* and *A. venetianus*^32,33^, and just reported in UTI isolates of *K. pneumoniae*^34^. TEM-imaging of DA14734 *wzc*^P642S^ showed an increased amount of capsule, distortions in the cell envelope, less defined LPS, and apparent shedding of the capsule from the cell surface (Fig. 4e). We also observed an increase in electron density in the periplasm, which could suggest accumulation of partly- or differently synthesized CPS components, as previously observed for mutations that block capsule synthesis at specific steps^35^. Overall, our findings indicate that the *wzc*-mediated hypermucoidy is not a simple overproduction of capsule, and it translates into biofilm phenotypes that differ from those in other types of hypermucoid isolates, like deletion in the capsule regulator *yrfF* we have observed earlier^18^.

Hypermucoviscosity itself was not a prerequisite for increased biofilm capacity in the evolved mutants. IA565 *rcsD*^G217R^ maintained the parental-like morphotype but displayed one of the most robust biofilms, especially on silicone with/without fibrinogen (Fig. 4c). RcsD is a histidine phosphotransferase in the Rcs system involved in the envelope stress response and positive capsule regulation. Balanced expression of Rcs components is required for biofilm formation and infection in the host^36,37^. This mutation in the periplasmic region of RcsD is likely a loss-of-function as the constructed gene deletion also led to increased biofilm formation without affecting the morphotype (Fig. 4b). In contrast to *wzc* mutants, loss of capsule mutants, especially when combined with fimbrial adhesin mutations, formed biofilm on polystyrene. Loss of capsule also changed how the biofilm was arranged on the silicone peg surface: instead of continuously covering the silicone, individual clusters of cells formed, suggesting increased aggregation between the cells in contrast to interaction with the surface (Fig. 4e). Taken together, these results illustrate that various levels and forms of capsule production can be advantageous in biofilms, but the biofilm properties can vary dramatically and depend on the specific mutation, strain background, and surface.

### Structural variations in the lectin domain of MrkD affect initial attachment, biofilm structure, and surface preference

Type 3 fimbriae mediate attachment of *K. pneumoniae* to catheter-related surfaces^38,39^, and mutations in type 3 fimbriae-related genes (*mrkD, mrkJ, yfiN, yfiR*) were commonly selected as better biofilm formers (Fig 2d and Fig 3). The type 3 fimbrial adhesin MrkD has been predicted to resemble the type 1 fimbrial adhesin FimH with a two-domain structure: a lectin domain (amino acid residues 24-184) with a binding pocket and a fimbria-anchoring pilin domain (amino acid residues 185-332)^40^. All our selected MrkD mutations mapped to the lectin domain that is responsible for receptor binding and reverts into a high-affinity conformation^40^ like FimH^41^ (Fig. 5a). Two of the genetic changes (T59C and in-frame deletion Δ(G40-V43)) overlapped across the strains; however, the selection of diverse *mrkD* variants was most common in IA565, where different *mrkD* alleles co-existed in the same evolving population or were combined in the same clone (Fig. 5b).

**Fig. 5.**
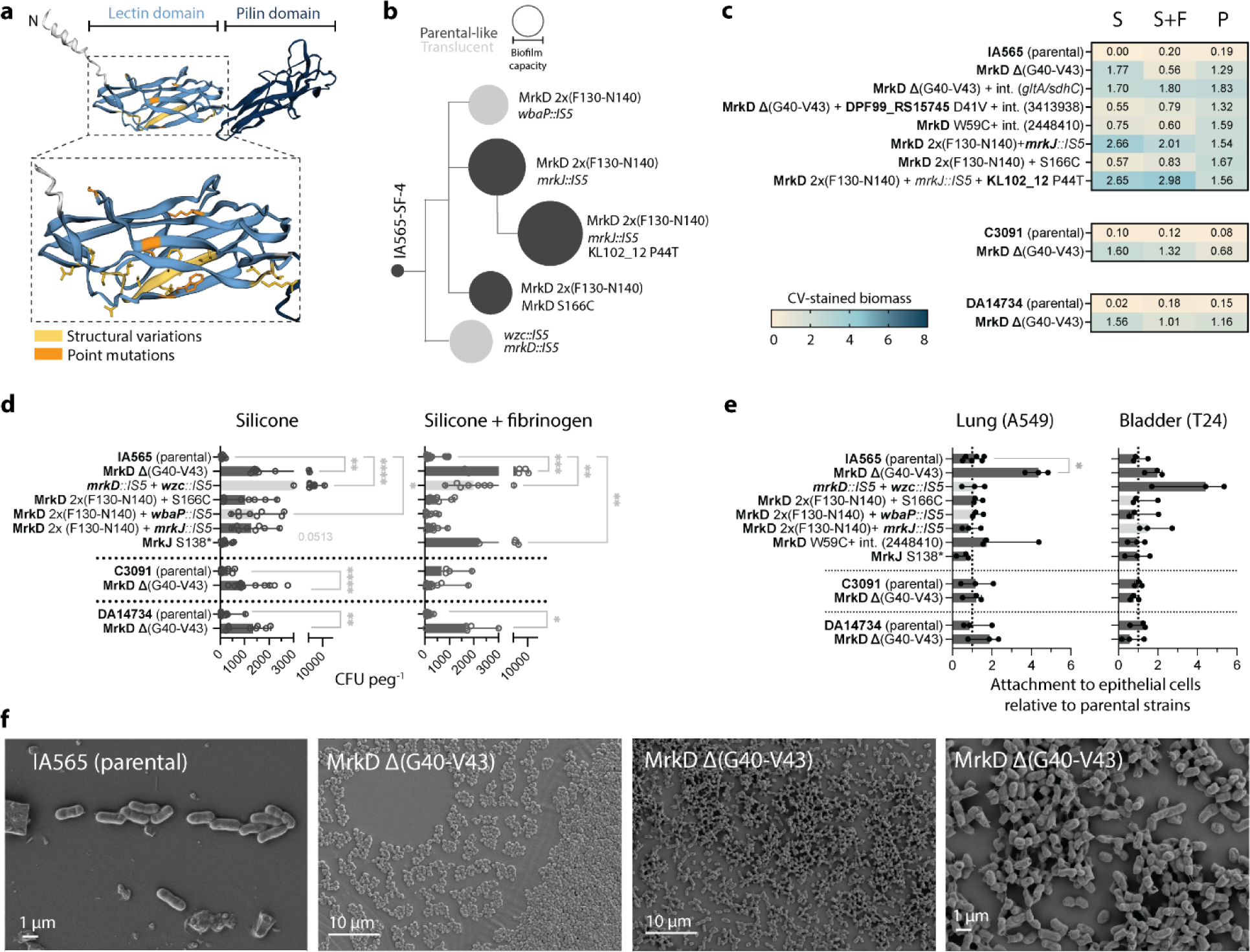
Characteristics of selected structural type 3 fimbriae mutants. **a,** Predicted MrkD protein structure (PDB K7R402) with marked mutated positions, **b,** Evolutionary trajectories in the IA565-SF-4 lineage. Biofilm capacity refers to the conditions during evolution (silicone with fibrinogen), **c,** CV-stained biofilm biomass on silicone pegs, silicone + fibrinogen pegs, and polystyrene wells after 48 h growth in evolved *mrkD* mutants. Heat map values illustrate medians of at least four biological replicates, **d,** Attachment to pegs after 3 h of growth. Results illustrate eight biological replicates with medians and 95% Cl. Comparisons between parental strains and mutants were made by the Kruskal-Wallis test, followed by Dunn’s multiple comparison test for IA565 and the Mann-Whitney test for C3091 and DA14734, with significance at ρ<0.05. **e**, Attachment of exponentially grown bacteria to A549 lung and T24 bladder epithelial cells. Bars show a median of three independent experiments with two technical replicates. Statistical significance was assessed by one-way ANOVA followed by Dunnett’s multiple comparison test; results were considered significant when p<0.05. **f**, SEM images of parental C3091 (left) and *mrkD* mutants (middle and right) grown for 16 h on silicone-coated pegs.

In contrast to *wzc* mutants, structural changes in the MrkD adhesin increased biofilm formation on polystyrene, reflecting altered interaction with different surfaces (Fig. 5c). Capsule loss mutants were often selected together with type 3 fimbria/c-di-GMP changes (e.g., *mrkD*, *mrkJ*::*IS5*), and such IA565 clones displayed the highest increase in biomass on all surfaces. Since fimbrial adhesins mediate the initial attachment of *K. pneumoniae* to abiotic surfaces^40,42,43^, we assessed the early attachment to pegs with or without fibrinogen (Fig. 5d). Early attachment to silicone increased 10 to 50-fold relative to the parental strains. A capsule-loss mutant with an IS5 insertion in *mrkD,* eliminating the tip adhesin, was particularly good at early attachment to silicone pegs with up to 50-fold higher CFU after 3 h. Although such a change might seem counterintuitive, MrkD is dispensable for biofilm formation on abiotic surfaces as long as MrkA is still present^44^. Mutants carrying a ten amino acid duplication (F130-N140) in *mrkD* were better at binding exclusively to silicone. The *mrkD*^Δ(G40-V43)^ deletion also improved attachment to silicone in all strains, although the most substantial effect was seen for IA565, especially with fibrinogen. SEM images showed that *mrkD*^Δ(G40-V43)^ was more likely to form cell clusters than a continuous monolayer at the early stages (16 h) of biofilm formation (Fig. 5f), reminiscent of the capsule-loss mutants (Fig. 4d).

To study mutational effects on the interaction with a biological surface, we assessed attachment to lung and bladder epithelial cells (Fig. 5e). The frequently selected *mrkD*^Δ(G40-V43)^ showed increased attachment to lung epithelial cells within 30 min but only in the respiratory isolate IA565. The same mutation in C3091 or DA14734, which carry a different *mrkD* allele, did not change the attachment to lung or bladder epithelial cells, suggesting a strain-specific behavior. The *mrkD*^W59C^ variant was trending towards an increase on lung cells and the no capsule mutant with *mrkD*::*IS5* was attaching better to bladder cells. Thus, the strong biofilm-forming MrkD variants have highly specific phenotypes whose fate would differ in niches with different abiotic surfaces and cell types.

### Regulation of type 3 fimbriae and cellulose production is interconnected in c-di-GMP-related mutants

Production of fimbrial adhesins and polysaccharides is regulated by c-di-GMP in many bacterial species^45–47^. Regulation of the type 3 fimbriae *mrk* operon in *K. pneumoniae* is the most well-characterized example of c-di-GMP dependent regulation in this pathogen^48–51^. The pool of c-di-GMP, essential for the activity of the MrkH transcriptional regulator, is maintained by the opposing activities of the MrkJ phosphodiesterase and the YfiN diguanylate cyclase, controlled by the repressor protein YfiR (Fig. 6a). Seemingly inactivating mutations in *mrkJ* were especially common in IA565, while changes in *yfiN* and *yfiR* were observed in C3091 (Fig. 3). Three uncharacterized EAL/GGDEF proteins were also targeted (Supplementary Fig. 3), including the DL426_09630^A203V^ wrinkly mutant, and we constructed in-frame deletions of these genes to determine the effects of the mutations. We reasoned that mutants with better biofilm capacity would overproduce type 3 fimbriae due to higher c-di-GMP levels and increased *mrkA* expression. Indeed, most mutations increased the *mrkA* expression, sometimes more than 100-fold (Fig. 6b). However, the expression was mutation-, strain-, and growth state-dependent. Interestingly, the C3091 *mrkJ*^A69P^ mutant had an increased *mrkA* expression in the exponential phase but a more than 5-fold decreased expression in stationary cultures. Similarly, but with opposite effects, C3091 *yfiR*^V78G^ had a >35-fold decreased *mrkA* expression in exponential, but a 20-fold increase in stationary phase, which might be the result of misregulation of c-di-GMP turnover via released repression on YfiN^52^. In comparison, the within-host evolved isolates from the outbreak with mutations in c-di-GMP-related genes had a remarkably high (up to >300-fold) increase in *mrkA* expression, especially in stationary phase.

**Fig. 6.**
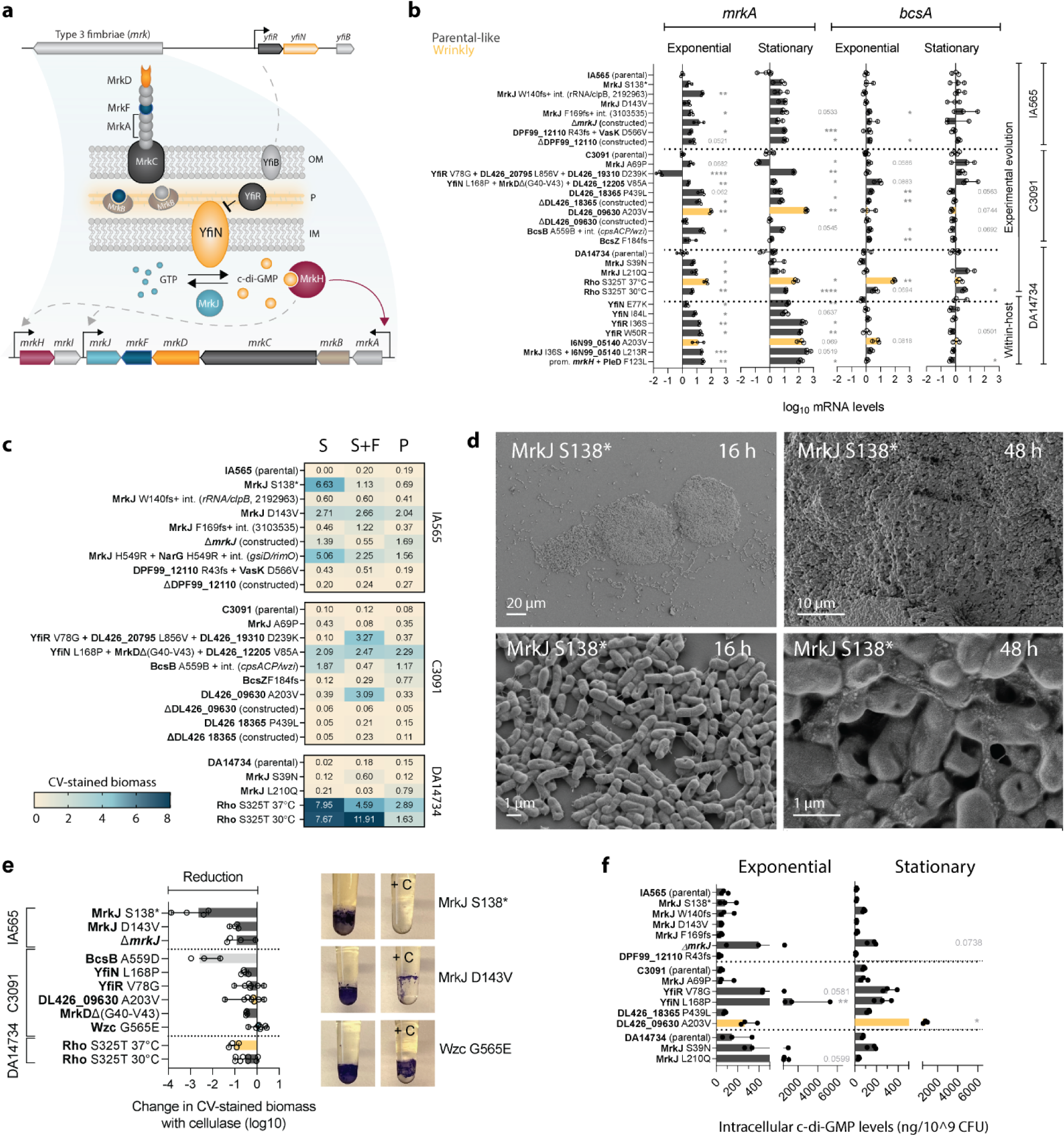
Interconnection between type 3 fimbriae, cellulose, and c-di-GMP in biofilm- and within-host evolved mutants. **a,** Schematic illustration of the type 3 fimbriae operon organization and expression regulation, **b,** *mrkA* and *bcsA* mRNA levels in selected mutants during exponential and early stationary growth normalized to the respective parental strain. Results from three biological replicates with 95% Cl are shown. Comparison between parental strains and mutants was done by one-way ANOVA, followed by an unpaired t-test with Welch’s correction. Mutations shown for within-host evolved isolates (DA14734) refer to mutations responsible for the biofilm phenotype, but the isolates contain additional changes^18^, **c,** CV-stained biofilm biomass on silicone pegs, silicone + fibrinogen pegs, and in polystyrene wells after 48 h growth. Results from at least four biological replicates, **d,** SEM images of *mrkJ*^S138*^ (IA565) on silicone pegs after 16h and 48h of growth, **e,** Effect of cellulase treatment on biofilms growing on silicone-coated pegs for 48 h. Results from eight biological replicates with 95% Cl are shown. Images show the pegs with (+C) and without cellulase. **f,** Quantification of c-di-GMP by LC-MS in exponential and early stationary cultures. Results from three biological replicates with 95% Cl are shown. Comparison between parental strain and mutants was done by Kruskal-Wallis test, followed by Dunn’s multiple comparison test. Differences are considered statistically significant when p<0.05. * p<0.05, ** p<0.0l, *** p<0.001,**** p<0.0001.

From this, it is clear that differences in *mrkA* expression alone cannot fully explain the biofilm capacity in *mrkJ*, *yfiN/R*, and other EAL/GGDEF protein mutants (Fig. 6c), illustrating that a combination of factors influences the phenotype. SEM imaging of the IA565 *mrkJ*^S138*^ mutant with the strongest biofilm phenotype revealed that while initially, the biofilm topology resembles that of loss-of-capsule with 80-100 µm cell clusters, at later time points, it develops multilayer biofilms with prominent ECM (Fig. 6d). YfiN/R has been connected to expression of the *bcs* cellulose operon^51^ and we also found changes in *bcsB*, which encodes a co-catalytic membrane protein in c-di-GMP-dependent synthesis and transport of cellulose, and the BcsZ endoglucanase, hydrolyzing cellulose. Given that cellulose regulation in *K. pneumoniae* biofilms is poorly understood, we determined both the mRNA levels of *bcsA* and the effect of cellulase (1,4-endoglucanase) on growing biofilms. While the *bcsA* expression was not affected in most mutants (Fig. 6b), biofilm formation in specific EAL/GGDEF protein and cellulose synthesis mutants (*mrkJ*^S138*^, *mrkJ*^D143V^, *bcsB*^A559D^) was eliminated by cellulase (Fig. 6e). In other cases, the reduction in biomass was more moderate (Supplementary Fig. 4). Biofilms of hypermucoid Wzc mutants were not affected at all, illustrating a different biofilm composition (Fig. 6e). Taken together, these results suggest that cellulose is a crucial ECM component in specific evolved *K. pneumoniae* mutants and that the c-di-GMP dependent regulation is not transcriptionally mediated, like for type 3 fimbriae, but acts at a posttranslational level. This observation is in agreement with a reported direct interaction between c-di-GMP and the PilZ domain of BcsA in other bacteria^45,53,54^. Despite displaying changes in c-di-GMP-dependent processes, most of the studied mutants did not show any increase in global intracellular c-di-GMP levels as determined by LC-MS (Fig. 6f). This suggests that local or temporal concentration changes are the drivers in the c-di-GMP regulated pathways in these mutants, or more subtle changes not visible from bulk *ex vivo* measurements.

Since the *rho*^S325T^ mutant also displayed a wrinkly morphotype (Fig. 1b), we analyzed its effects on *mrkA* and *bcsA* expression and cellulose production. This mutant showed an extremely robust biofilm formation (Fig. 6c), even at 30℃, where the colony morphotype reverted to parental, and was among the mutants with the highest increase in both *mrkA* and *bcsA* expression (Fig. 6b). The biofilm was also sensitive to cellulase, suggesting that cellulose is a crucial biofilm component. However, unlike the *mrkJ* mutants, *rho*^S325T^ increased *bcsA* mRNA levels >80-fold, suggesting an alternative regulatory pathway that is probably not directly linked to c-di-GMP. The S325T mutation is near one of the positions (324, in the R-loop) found to be crucial for transcriptional termination^55^.

### Acute virulence can be lost as a trade-off to increased biofilm capacity

Although the *K. pneumoniae* strains were cycled without any host material/defense (except for fibrinogen), the mutated structures and processes are important in pathogenesis, and the changes can be predicted to alter the virulence of the bacteria. The complement system is one of the first-line defenses against pathogens in the human host, and serum resistance of a bacterium can help predict the potential for systemic infection^56^. As expected, loss-of-capsule mutants were sensitive to human serum; however, despite the predicted serum resistance due to hypermucoidy, *wzc* mutants were extremely serum sensitive (>5 log increased killing within three hours) (Fig. 7a). An equally high serum sensitivity was observed among outbreak hypermucoid *wzc* isolates^18^. Additional mutations in *prc*, which is important in complement evasion in *E. coli*^57^, did not rescue serum sensitivity in *wzc* mutants. However, this sensitivity was abolished if the mutants were grown as a biofilm first and then exposed to serum (Fig. 7b). In contrast, mutational change or constructed knock-out of *rcsD* did not affect serum sensitivity, which agrees with the parental-like morphotype. Interestingly, in C3091, *mrkD*^Δ(G40-V43)^ and *mrkJ*^A69P^ showed better survival in human serum. This illustrates that in addition to LPS and capsule, other cell surface structures or interactions between them can be crucial in protecting the cells from complement attacks by limiting the activation or deposition of complement proteins on the cell surface in a strain-dependent manner^58,59^.

**Fig. 7.**
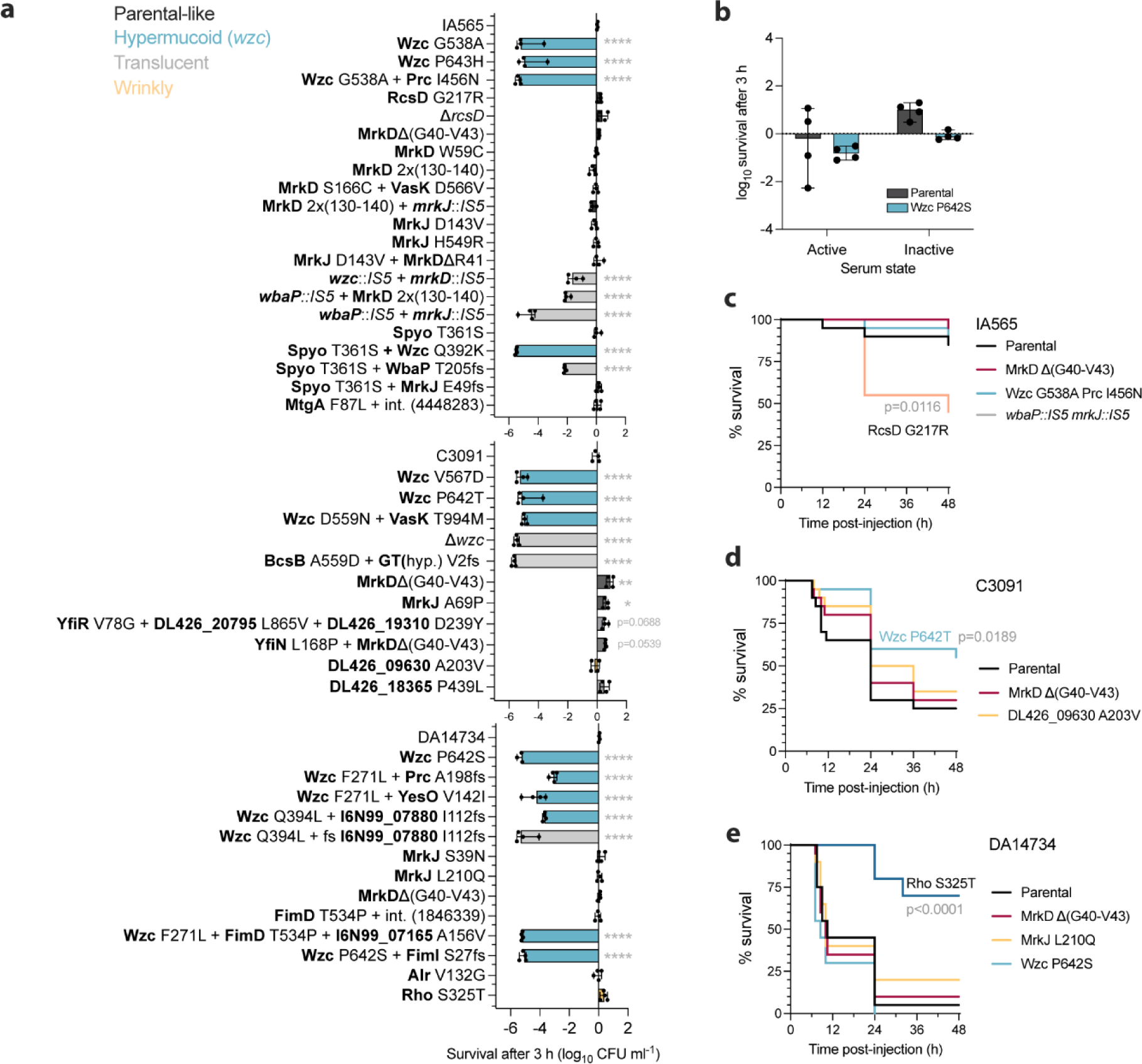
Virulence features in evolved mutants. **a,** Serum killing assays with planktonic cultures as Iog10 difference in CFU/ml after 3h exposure to human serum. Bars show medians of four biological replicates. Comparisons to parental strains were done by one-way ANOVA followed by Dunnett’s T3 multiple comparison test; significant comparisons are shown p<0.05. **b,** Serum killing assay with biofilms of the parental strain DA14734 and *wzc*^P242S^ mutant. The biofilms were grown for 48 h on silicone-coated pegs and exposed to human serum for 3 h. Survival of *G. mellonella* larvae infected with mutants (10^5^ CFU) derived from **c** IA565, **d** C3091 and **e** DAI 4734 strains. Pairwise comparisons of mutants survival curves (n=20 larvae) against parental were done by log-rank (Mantel-Cox) test, p values for significantly different comparisons are shown (p<0.05). * p<0.05, ** p<0.01, *** p<0.001, **** p<0.0001.

To analyze the pathogenic potential of biofilm mutants in a more complex system, we assessed the survival of a set of mutants, including capsule, fimbriae, or c-di-GMP signaling-related changes in *Galleria mellonella* larvae (Fig. 7c-e). This infection model has been successfully used for multiple species, including opportunistic *K. pneumoniae*^60–62^. Some mutations had a strain-dependent effect on virulence. For example, *wzc* point mutations at amino acid position P642/643, which led to high sensitivity in human serum in all strains, significantly decreased the killing of larvae in C3091 but not in IA565 or DA14734. For DA14734 *wzc*^P642S^, there was a tendency towards faster killing (Fig. 7e), in contrast to other *wzc* mutants from the outbreak^18^. The strong biofilm-forming mutant IA565 *rcsD*^G217R^ had increased larvae killing, consistent with its resistance to human serum (Fig. 7c). In contrast, the particularly strong biofilm-forming mutant DA14734 *rho*^S325T^ was partially attenuated in larvae, indicating that biofilm capacity is not automatically coupled to increased killing.

## Discussion

Since interaction with surfaces is often a central part of nosocomial *K. pneumoniae* infections^6,7,63,64^, exploring the adaptive changes that lead to altered biofilm phenotypes can give important insights into the pathoadaptivity of this bacterium. Here, we demonstrate that *K. pneumoniae* can rapidly evolve improved biofilm formation when subjected to cycling *in vitro* on abiotic surfaces, including those mimicking the typical catheter surface. In a clinical situation, this could translate into the selection of bacterial subpopulations that initiate infection of intubated or catheterized patients due to altered adhesion abilities. By comparing the *in vitro* evolution of the outbreak clone DA14734 to changes identified in patient isolates^18^, we show that short-term experimental evolution can replicate some of the most common adaptations selected within the host and possibly explain why they were established during infection. The morphotypes we identify as advantageous in biofilm, for example, hypermucoid and translucent, have also been linked to globally transmitted high-risk *K. pneumoniae*^28^. A rarely reported wrinkly morphotype was recently connected to adaptation in carbapenem-resistant hypervirulent *K. pneumoniae*^65^ and is generally associated with chronic infections in other bacteria^12,47,66,67^.

The evolutionary trajectories of the three opportunistic strains isolated from UTI, wound, and trachea exhibited strong genetic parallelism, where vital cellular structures and processes were the main mutational targets. While we see strong convergent evolution across strains, there seems to be a strain-specific bias in what changes are selected the most and how strong the phenotypic effects are. This illustrates that while certain shared factors are essential for biofilm formation in *K. pneumoniae*, their contributions can vary depending on the genetic background, possibly diversifying the adaptability to specific niches. Strain-dependent selection bias was apparent on the target level (capsule vs. fimbrial mutations), on the gene level (e.g., *mrkJ* vs. *yfiN/yfiR*), and on biofilm phenotype (*wzc* mutants in IA565 vs. C3091/DA14734). Missense mutations in *wzc* leading to a hypermucoid phenotype were more often selected in C3091 and DA14734, and the phenotype was much more prominent in these strains. Differences in capsule-mediated phenotypes are likely due to different capsule types, and strain influences on capsule involvement in *K. pneumoniae* biofilms have also been observed by others^68^. Similarly, mutations affecting the regulation of type 3 fimbriae and the structural component MrkD were often co-selected with mutations abolishing capsule production, similar to a recent study^69^. Such mutants likely expose fimbriae more and were primarily found in the IA565 background. This could point to a higher evolvability of the *mrkD-*allele in IA565, which differs from the most abundant and stronger binding allele present in the other two parental strains^40^. IA565 also carries an additional *mrkD_1P_* on a plasmid^70^, although we found no mutations in that locus. Differences in specific c-di-GMP-related changes, such as different levels of fimbrial expression or cellulose production, are likely attributed to different networks of EAL/GGDEF domain proteins in specific strains that drive particular responses, as complex c-di-GMP signaling networks show extensive variation even within species^71–73^. Considering the vast genetic diversity of *K. pneumoniae*^8^, it is essential to study such strain dependencies systematically and identify the key factors influencing the response in different genetic backgrounds.

Our results here point to biofilm growth as a selective pressure for the emergence of both the hypermucoid and capsule-loss mutants that we also identified as within-host-selected infection site adaptations during the Uppsala hospital outbreak^18^. In parallel to our work, another study reported the selection of *wzc* mutants in structured environments, which they achieved by serial passaging in 24-well plates^74^. Widespread alteration or loss of capsule production via *wzc* and *wbaP* mutations was also recently reported among globally disseminated *K. pneumoniae* ST258 isolates^28^. However, this study suggested that *wbaP* mutants in ST258 form surface-attached biofilms, while *wzc* mutants do not^28^. Importantly, we find that *wzc* mutants strongly attach to silicone or fibrinogen, but not to polystyrene microtiter plates, which was the method used by Ernst et al.^28^. On the other hand, we see apparent phenotypic differences of these mutants in different strain backgrounds, which could be another explanation for the discrepancies between the studies. Additionally, in contrast to mutants that carry only *wzc* mutations, within-host selected isolates with additional genetic changes did not form biofilm in the same conditions^18^. This could be due to additional mutations in other genes that possibly compensate for specific trade-offs of the *wzc* phenotype, although this remains to be determined. Biofilm growth rescues *wzc* mutants from the extreme planktonic serum sensitivity we otherwise observe when *wzc* mutations are present alone or with other genetic changes. Similarly, LPS mutations improving biofilm growth in *P. aeruginosa* can make planktonically growing cells more sensitive to antimicrobial agents^75^. Such trade-offs between planktonic and biofilm growth affecting the survival in the host could generate a subsequent selection for compensatory mutations.

The growing number of studies identifying Wzc as a mutational target linked to infections warrants a deeper understanding of its role in the observed phenotypes. Mechanistically, there are multiple possibilities for how *wzc* mutations could affect the protein function and, consequently, the entire capsule synthesis machinery. For example, alterations in the interaction with the Wza transporter on the periplasmic side could directly affect the export of components from the periplasm. The increased periplasmic density we observe with TEM could support this. Alternatively, changed interactions with the Wzc phosphatase in the cytoplasm could interfere with the cycling between the phosphorylated and dephosphorylated states and, thus, the oligomerization of Wzc itself. Altered phosphorylation levels have just been reported in *wzc* mutants from UTI *K. pneumoniae*^34^. The key phenotypic feature that we observed both in single *wzc* missense mutants and the combined *wzc* mutants from the outbreak is the extreme sensitivity to serum in planktonic conditions. Based on our TEM observations, we propose that this is due to multifactorial effects on the cell envelope, including increased shedding of high molecular weight capsular polysaccharides. To further support this, a weakened association of the capsule with the cell surface in such *K. pneumoniae* mutants has just been reported^34^. In addition, the integrity of the whole cell envelope seems to be disturbed, possibly due to sequestration of undecaprenyl phosphate and the accumulation of partly synthesized components^35^. Frameshift mutations in the Prc protease (processing of PBP3), deletion of which results in leakage of periplasmic proteins in *E. coli*^76^, always coincided with *wzc* mutations in our study, raising the question about a possible interplay between these changes. Inactivation of *prc* via transposon insertion also increases capsule production in *K. pneumoniae*^77,78^.

The biofilm-conferring mutations identified here affect factors important for *K. pneumoniae* pathogenesis, for example, capsule or type 3 fimbriae, suggesting a possibly altered virulence status. However, virulence profiles in *G. mellonella* larvae did not directly correlate with biofilm capacity, meaning that excellent biofilm-forming strains can have reduced (e.g., *rho*), increased (e.g., *rcsD*), or non-changed acute virulence. Thus, the relationship between biofilm formation and acute/systemic virulence depends on the exact genetic change underlying the biofilm phenotype, which is in agreement with our recent observations in within-host evolved isolates^18^. Similarly, a study of *K. pneumoniae* defective in biofilm formation found no direct correlation between loss of biofilm formation and the ability to cause lung infections in mice^79^. However, we have associated the high biofilm-forming isolates with a gastrointestinal colonization advantage in mice^18^. These results illustrate the complex interplay between genetic variations in mediating the overall phenotype and the within-host selection of bacteria with altered capacity for infection through indwelling medical devices.

## Materials and Methods

### Bacterial strains and growth conditions

Three clinical strains of *K. pneumoniae* were used: (i) IA565 (DA11912, ST105), a clinical tracheal aspirate isolate, originally from the University of Iowa Hospitals and Clinics Special Microbiology Laboratory^26^; (ii) C3091 (DA12090, ST14), a *K. pneumoniae* UTI isolate from Walter Reed Army Medical Center^25^; and (iii) DA14734 (ST16), an ESBL-producing *K. pneumoniae* that was the index isolate from an outbreak at Uppsala University Hospital^23^. Clone and population strain numbers from the evolved biofilm lineages with information on all mutations are noted in Supplementary Table 1. Bacteria were grown in Brain Heart Infusion broth (BHI, Oxoid) both before inoculation and for biofilm growth. For the preparation of electrocompetent cells and serum-killing assays, planktonic bacteria were grown in Lysogeny Broth (LB, Sigma). Antibiotics used to select transformants during strain construction were cefotaxime (10 mg/L, Sigma) and kanamycin (50 mg/L, Sigma).

### Setup for experimental evolution on pegs

The cycling of *K. pneumoniae* IA565, C3091, and DA14734 was performed on a 3D printed FlexiPeg device^19^ with high-temperature resin (HT) pegs (Form 2 3D-printer) and silicone (polydimethylsiloxane, PDMS)-coated pegs. Silicone-coated pegs were further coated with 100 mg/L fibrinogen as described before^18^ when indicated. Overnight (O/N) cultures of 10 independent lineages for each strain on HT pegs and 6 lineages for each strain on silicone-coated pegs and silicone-coated pegs with additional fibrinogen coating were diluted 100-fold or 10.000-fold and 150 µl transferred to a 96-well microtiter plate (NuncTM) where the peg lid was then inserted. The biofilms formed on the pegs were transferred to fresh medium after 24 h and were harvested after 48 h by vortexing for 2 min as previously described^19^. We first performed a pilot cycling on high-temperature resin (HT) pegs as a proof of concept. After each 48h cycle, 0.25% or 25% of the disrupted biofilm population was transferred to initiate biofilms on new pegs. No apparent differences were observed in the rate or range of mutations selected with the different bottlenecks, and since the parental strains did not reach as high CFU/peg with the smaller bottleneck, which led to an increased population loss rate, cycling was only continued with the 25% bottleneck in the following experiments. Cycling was continued for 5 cycles (non-coated BHI conditions) or 6 cycles (silicone with/without fibrinogen, BHI).

### Biofilm growth on pegs

The biofilms were grown in the FlexiPeg device for 12, 16, 20, or 48 h according to the general protocol^19^. The biofilms were harvested by vortexing for 2 min or stained with 0.1% crystal violet (CV, Sigma) at different time points as previously described^19^.

### Biofilm formation in polystyrene microtiter plates

O/N cultures were diluted 10000-fold in BHI and 150 μl inoculated into a round bottom 96-well plate (Nunc) to initiate biofilm formation in microtiter plates. The plate was statically incubated for 48 h at 37°C. After 48 h, unattached cells were removed from the wells, and the plate was washed 3x with 1x sterile PBS, dried for 30 min at 37°C, and 180 μl of 0.1% CV (Sigma) was added. After 20 min of incubation at room temperature, the plate was extensively washed (3-4x) with PBS to remove any non-bound CV stain, dried for 20 min, and 180 μl of 10% acetic acid was added to solubilize the stain. The level of CV was quantified by measuring the absorbance at 540 nm with a Multiskan^TM^ FC Microplate Photometer (Thermo Scientific). The average value from the blank (acetic acid only) was subtracted from the sample values.

### Biofilm formation in the presence of cellulase

An aqueous solution of cellulase from *Aspergillus niger* (Sigma, product number C2065) was diluted 1:60 in BHI to yield approximately 20 mg/ml, and 150 µl were transferred to a flat-bottom 96-well plate (Nunc). O/N cultures were diluted 1:100 in BHI, and 1.5 µl were transferred to a 96-well plate with BHI + cellulase or BHI only, and the FlexiPeg lid was inserted into the plates. The biofilms were allowed to form for 24 h on silicone-coated pegs, then they were harvested by vortexing for 2 min and plated on LA plates for CFU counts.

### Exponential growth rate measurements

O/N cultures were diluted 1000-fold in BHI, and 300 μl were transferred to 100-well honeycomb plates. Four to five biological replicates were used for each isolate. The plate with cultures was incubated in a BioScreen C (Oy Growth Curves Ad Ltd) for 16 h at 37°C with shaking. OD_600_ measurements were taken every 4 min. Results were analyzed using the R script-based tool BAT 2.0^80^ to determine relative growth rates.

### Strain construction by λ Red recombineering

In-frame deletions of genes mutated during the cycling in the biofilm were constructed in IA565 and C3091 background. Strains were first transformed with the pSIM5-CTX and selected on LB agar with cefotaxime (10 mg/L) at 30°C. pSIM5-CTX carries the temperature-inducible λ Red recombineering system and is derived from the original pSIM5^81^. A *kan-sacB-T0* cassette was amplified by PCR using Phusion High-Fidelity DNA Polymerase (Thermo Fisher Scientific Inc.) with primers including 40 bp flanking homology regions to the gene of interest. The cultures were grown until early exponential phase (OD_600_ ∼0.3) and the λ Red system was induced by 15 min incubation in a 42°C shaking water bath. The cells were made electrocompetent by washing 4-5 times in sterile 10% glycerol and 100-500 ng of the purified *kan-sacB-T0* cassette was electroporated. The cells were recovered in LB or BHI overnight at 30°C. Transformants were selected on LB agar with kanamycin (50 mg/L) at 30°C and checked for sucrose sensitivity on 5% sucrose plates and the carriage of the pSIM5 plasmid on LB agar with cefotaxime (10 mg/L). Successful transformants were used for another λ Red recombineering step with a linear ssDNA fragment containing 40 bp homologous regions directly upstream and 40 bp downstream of the gene of interest to delete the *kan-sacB* cassette. Transformants were selected on 5% sucrose and PCR-verified for the correct deletion. Primers used for the construction can be found in Supplementary Table 2.

### Genomic DNA extraction, whole-genome sequencing (WGS), and bioinformatics

Genomic DNA was extracted from 500 µl of O/N cultures using the Epicentre MasterPure^TM^ DNA purification kit (Illumina Inc.) according to the manufacturer’s instructions. For population sequencing after evolution, we did a short pre-growth before DNA extraction to reach approximately 4×10^8^ CFU/ml for populations with low CFU (below 10^7^ CFU/peg) numbers. Depending on the lineage, the pre-growth resulted in 5 to 16 generations of growth, and a pilot run showed that this short pre-growth did not change the frequencies of morphotypes in the evolved population. The two parental strains, IA565 and C3091, were sequenced using the Pacific Biosciences II technology at the Science for Life laboratory sequencing facility and assembled to create reference genomes. For resequencing of mutants and correction of possible PacBio sequencing errors in the reference genomes, DNA samples were prepared according to Nextera© XT DNA Library Preparation Guide (Illumina Inc.), and sequencing was performed using a MiSeq^TM^ Desktop Sequencer (Illumina Inc.). The raw data was analyzed in CLC Genomic Workbench v.20 (Qiagen), where raw reads were trimmed and aligned to the reference sequences to analyze the genetic changes. Genetic changes were detected using the Microbial genomics module and resequencing SNP and InDel identification software in the CLC Genomics workbench. For clones, the cut-off for calling SNPs was set to >75%, and for InDels, a frequency of >0,5 among reads was used, and each change was manually inspected. Genetic changes with at least 20% frequency were included for populations unless lower-frequency mutations known to be present in clones were found. Protein sequences were further analyzed with BLASTp (NCBI) and a conserved domain database^82^ for functional determination. The complete genome sequences of IA565 (DA11912) and C3091 (DA12090) have been deposited under BioProject PRJNA473315 and PRJNA473316, respectively, and the outbreak index isolate under PRJNA857654. Sequence files for all mutants in this study have been deposited in the Short Read Archive (NCBI) under BioProject PRJNA1048869.

### Transmission electron microscopy

Bacterial strains were grown at 37°C with shaking (180 rpm) overnight in BHI and 2 ml centrifuged at 8000xg for 5 min. The supernatant was removed, 5 ml of ice-cold fixative solution (2.5% glutaraldehyde and 1% paraformaldehyde in 0.1 M sodium cacodylate buffer) was added, and the fixation continued overnight at 4°C. Further sample preparation and imaging were done at the BioVis EM node (Rudbeck’s laboratory, Uppsala University). The samples were post-fixed with 1% osmium tetraoxide in 0.1 M PIPES buffer and dehydrated in increasing ethanol concentrations (70%, 90%, and 100%). After dehydration, the samples were either negatively stained with uranyl acetate or further embedded in Agar 100 resin, ultrathin-sectioned (60 nm thickness), contrasted with uranyl acetate and lead citrate, and air-dried. The samples were imaged using a transmission electron microscope FEI Technai G2 at 80 kV.

### Scanning electron microscopy

The biofilms grown on silicone-coated pegs were fixed at 16 h or 48 h and prepared for SEM imaging as described before^19^. Images shown in Fig. 4d were colour-modified in Adobe Photoshop to distinguish the biofilm-covered area from the peg surface.

### Sedimentation assay

Bacteria were grown in BHI to late stationary phase, and cultures were centrifuged for 5 min at 1000x g. The sedimentation constant was calculated as the ratio of OD_600_ of the supernatant and the OD_600_ before centrifugation^77^.

### Extraction of capsular polysaccharides

Extraction of capsule from *K. pneumoniae* was done according to^77^. 100 μl of capsule extraction buffer (500 mM citric acid pH 2.0, 1% Zwittergent 3-10) was added to 0.5 ml of overnight cultures. After incubation at 50°C for 20 min, the mixtures were centrifuged for 5 min at 17,000xg at room temperature. Then, capsule components were precipitated by incubating 300 μl of supernatant with 1.5 ml of 99.5% ethanol at 4°C for 30 min. The precipitates were collected by centrifugation (20 min, 18,000 x *g*, 4°C) and air-dried.

### *mrkA* and *bcsA* mRNA quantification by qPCR

Bacteria were grown in BHI, and culture aliquots for RNA extraction were taken during mid-exponential (OD=0.5), late exponential (OD=1.0) and early stationary phase (6 h of growth). RNA was extracted using the RNeasy Mini Kit (Qiagen), and genomic DNA was removed using the TURBO DNA-free kit (Invitrogen). Complementary DNA (cDNA) was synthesized using the High-Capacity Reverse Transcriptase kit (Applied Biosystems) with approximately 500 ng of extracted RNA. The expression of *mrkA* and *bcsA* was assessed with qPCR (Illumina Eco Real-Time PCR System) using the SYBR Green Master mix (Thermo Fisher) relative to the expression of the *glnA* reference gene^83^. Primer sequences can be found in Supplementary Table 2. The mRNA levels were normalized to the respective parental strains mRNA levels for the respective conditions using the 2^−ΔΔ*C*^_T_ method^84^.

### c-di-GMP quantification by LC-MS

Selected clones were grown in BHI with shaking (180 rpm) at 37°C until mid-exponential phase (OD=0.5) or early stationary phase (6 h of growth), and culture aliquots (concentrated to OD 5) were washed three times in ice-cold sterile 1x PBS. The samples were kept at −80°C until the extraction was done at the Swedish Metabolomics Centre (Umeå, Sweden). Frozen samples were extracted using 400 µl 80% MeOH + salicylic acid-D6 (2 pg/µl). Supernatant (5 µl) from the centrifuged samples was injected into the instrument (Agilent 6490 triple quadrupole system). To make a calibration curve, c-di-GMP was serially diluted to produce a range from 31 fg/µl to 8 pg/µl. The amount of c-di-GMP was expressed as the mass of c-di-GMP (pg) per 10^9^ CFU in a sample.

### Survival in human serum

Selected clones were grown overnight in LB, diluted 10.000-fold in PBS, and approximately 10^5^ CFU mixed with human serum from male AB plasma (Sigma-Aldrich, product number H4522) in a total volume of 200 µl. Due to natural batch variation in human serum activity, the assays were performed either with undiluted serum or 40% dilution (in PBS) to achieve the same killing effect relative to the parental strains. The serum was heat-inactivated for 30 min at 56°C for negative control experiments. The CFU counts were performed at 0 h and after 3 h of static incubation at 37°C. For testing the biofilm’s sensitivity to human serum, biofilms were formed on FlexiPeg for 48 h, washed in PBS, and exposed to human serum for 3 h, after which CFU/peg was determined.

### Virulence in Galleria mellonella larvae

Selected mutants were tested for the killing of *Galleria mellonella* larvae. Larvae were purchased from Herpers choise (Uppsala, Sweden) and stored for a maximum of 3 days at room temperature before experiments. Only healthy larvae (no discoloration, active) of similar size (approximately 250-300 mg) were used for the experiments. Bacterial cultures grown overnight in BHI were diluted in PBS and 10 µl injected via the top right proleg using a 25 µl Hamilton 7000 syringe (Model 702 RN, 22S gauge needle). For each strain, 20 larvae (1 biological replicate per 10 larvae) were injected and monitored hourly for 7-12 h post injection on the first day, then at 24 h, 36 h, and 48 h post injection during incubation at 37°C. Aliquots from dilutions were plated for CFU counts to confirm the correct infectious load. Along with experimental larvae, 10 larvae were injected with 10 µl of sterile PBS to account for possible injection trauma, and 10 larvae served as non-injection control. The larvae were considered dead when not moving after stimulation with a pipette tip. In addition, larvae were given health index scores at 8 h, 12 h, 24 h, and 48 h post injection based on the previously described scoring system, considering melanization status and changes in movement/responsiveness^85^. A cumulative score was calculated for each replicate by adding the scores for each individual larva at different timepoints. Survival curves were analyzed by log-rank (Mantel-Cox) test in GraphPad Prism version 9.2.0 for macOS (GraphPad Software, San Diego, CA, USA).

### Mammalian cell culture

T24 bladder transitional carcinoma cells (ATCC HTB-4) were grown in McCoy’s 5a Medium Modified (Sigma) supplemented with 10% heat-inactivated fetal bovine serum (FBS). A549 lung carcinoma cells (ATCC CCL-185) were grown in RPMI 1640 (Gibco) containing 10% heat-inactivated FBS. Epithelial cells were seeded in tissue culture-treated 24-well plates 18-48 h prior to infection.

### Adherence to human epithelial cells

T24 and A549 epithelial cells were seeded densely enough to cover the entire bottom of the well and thereby limit bacterial binding to the plastic. Bacteria were grown in 2 ml of BHI broth overnight with shaking (180 rpm) at 37°C. The following day, cultures were used directly for infection with stationary phase bacteria or diluted 1:100 in BHI broth and grown with shaking at 37°C for 1.5 h for infection with exponentially growing bacteria. Each well was infected with 10^7^ bacteria. To synchronize infection, the bacteria were centrifuged onto the cells at 250xg for 5 min and left to adhere for 30 min at 37°C and 5% CO_2_. Cells were washed three times in PBS to remove unbound bacteria. The monolayers were solubilized in 0.2% sodium deoxycholate (DOC), and serial dilutions were plated on LA plates to enumerate the adhering bacteria.

## Author contributions

G.Z. and L.S. conceived the study and designed experiments. G.Z. performed experimental evolution, screening of clones, extracted genomic DNA from clones and populations, prepared samples for SEM and TEM imaging, c-di-GMP quantification, performed capsule extractions, sedimentation assays, *G. mellonella* infection assays, and strain constructions. G.Z. and P.C. measured biofilm capacity, early attachment to pegs, extracted RNA, and performed qPCR, serum killing assays, cellulase assays, and exponential growth rate measurements. M.W. performed experiments with epithelial cells. G.Z. and L.S. performed bioinformatic analysis. G.Z. and L.S. analyzed data and wrote the manuscript. G.Z. made the figures. All authors read, edited, and approved the final version of the manuscript.

## Acknowledgments

Parts of this work were presented at ASM Microbe 2019 (San Francisco, USA), EuroBiofilms 2019 (Glasgow, UK), and the 8^th^ National Infection Biology/Microbiology Meeting 2019 (Bålsta, Sweden). The authors would like to thank Victoria Sternhagen for performing the SEM analysis and data acquisition carried out within the Uppsala University academic cleanroom, a member of the MyFAb national research infrastructure. Monika Hodik from BioVis (Rudbeck Laboratory, Uppsala University) is acknowledged for her help with TEM sample preparation and imaging. The Swedish Metabolomics Centre (Umeå, Sweden) is acknowledged for the quantification of c-di-GMP by LC-QqQ-MS. Uppsala Antibiotic Center is acknowledged for funding of P.C. as a PhD student within the UAC research school. We also thank Dan I. Andersson and Vaughn S. Cooper for their comments on the manuscript draft.

## Supplementary videos

Supplementary videos are available on https://uppsala.box.com/s/u1vf7iha2kyxp0nztn85bt0sa03yrcq9

