## Supplementary Data 1 for "Rapid evolution of *Klebsiella pneumoniae* biofilms *in vitro* delineates adaptive changes selected during infection"

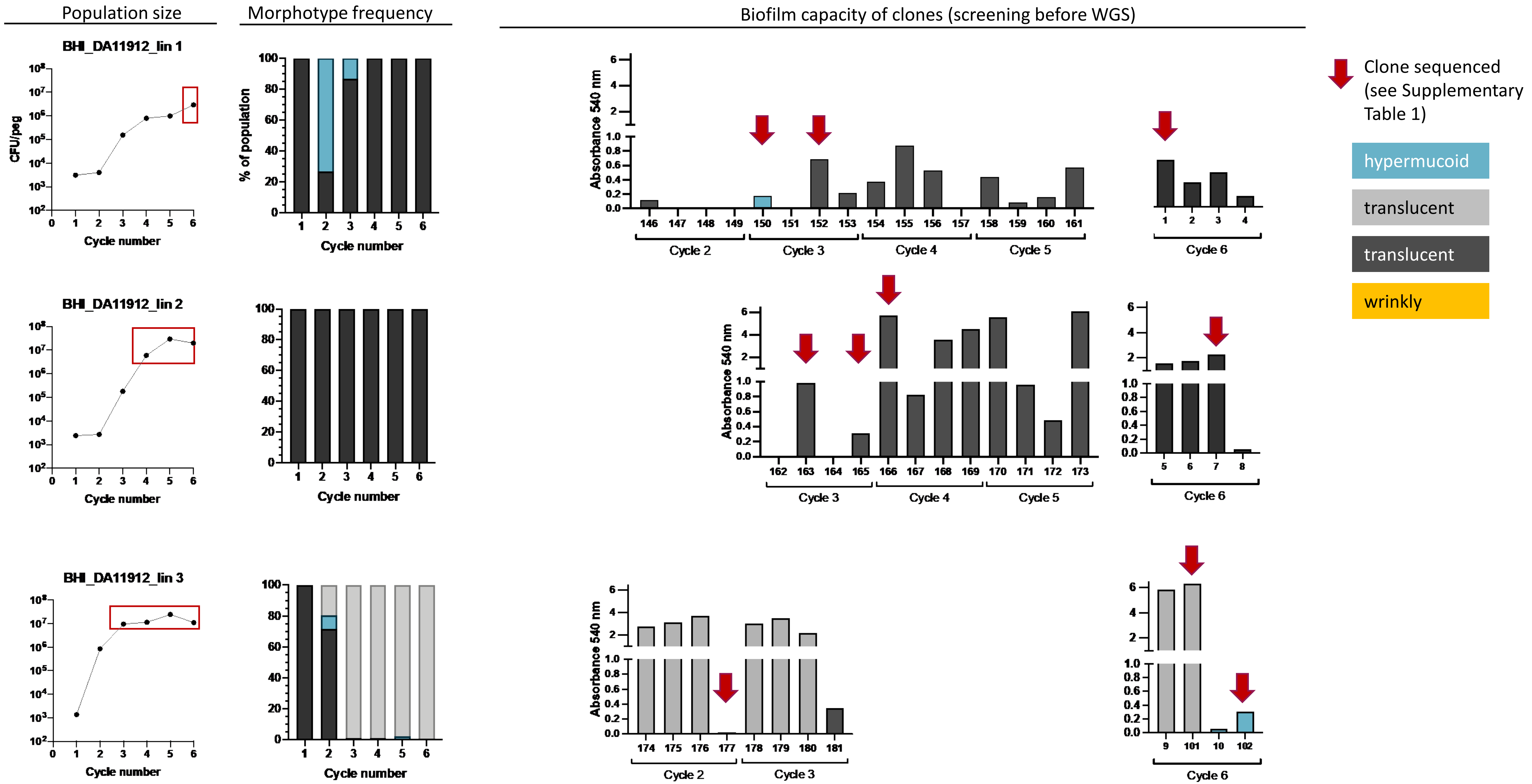
Population size
Morphotype frequency
Biofilm capacity of clones (screening before WGS)
BHI_DA11912_lin 1
BHI_DA11912_lin 2
BHI_DA11912_lin 3
CFU/peg
Cycle number
% of population
Absorbance 540 nm
Cycle 2
Cycle 3
Cycle 4
Cycle 5
Cycle 6
Clone sequenced (see Supplementary Table 1)
hypermucoid
translucent
translucent
wrinkly

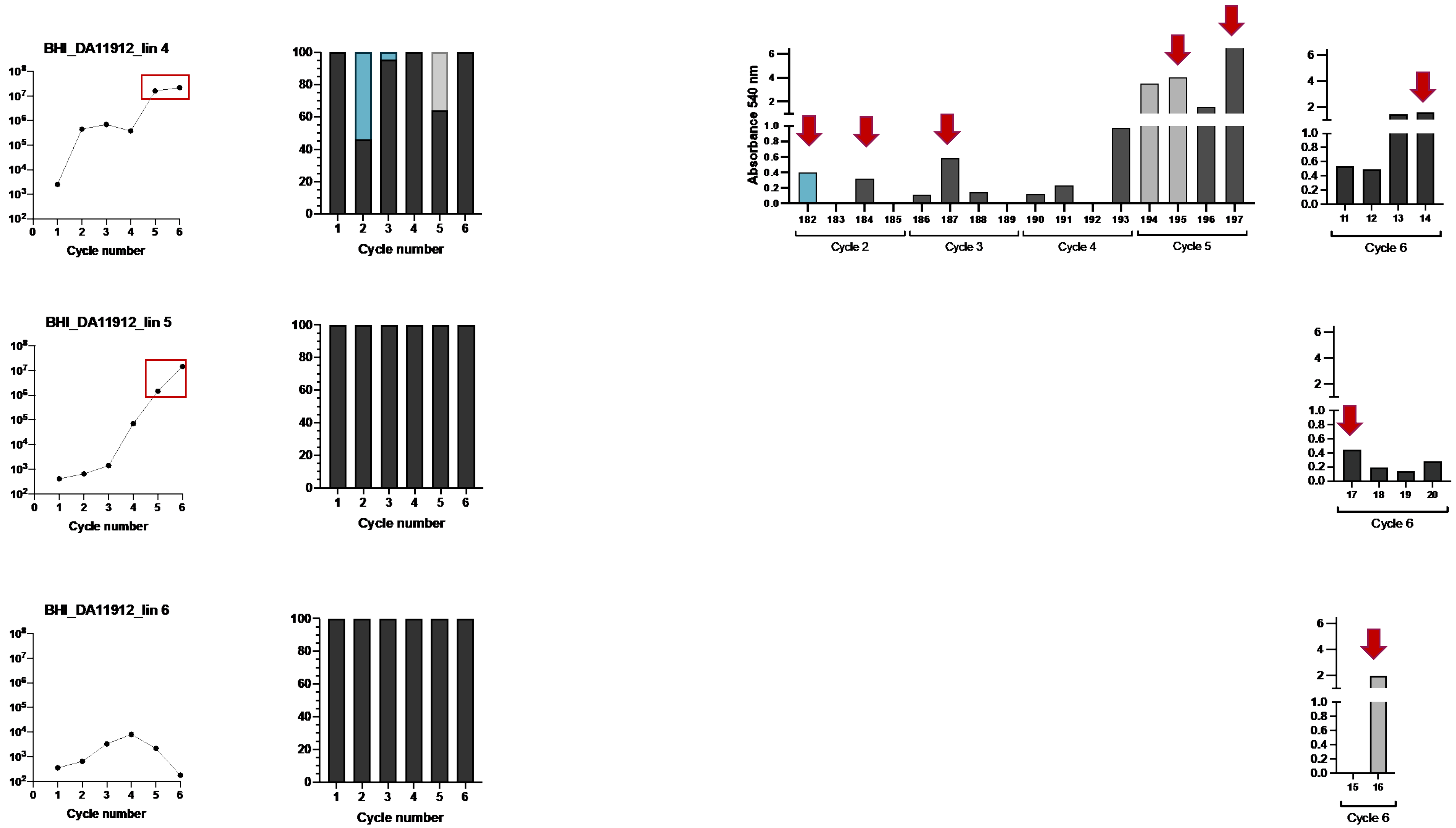

BHI_DA11912_lin 4
Cycle number
Absorbance 540 nm
Cycle 2
Cycle 3
Cycle 4
Cycle 5
Cycle 6
BHI_DA11912_lin 5
Cycle number
Cycle 6
BHI_DA11912_lin 6
Cycle number
Cycle 6

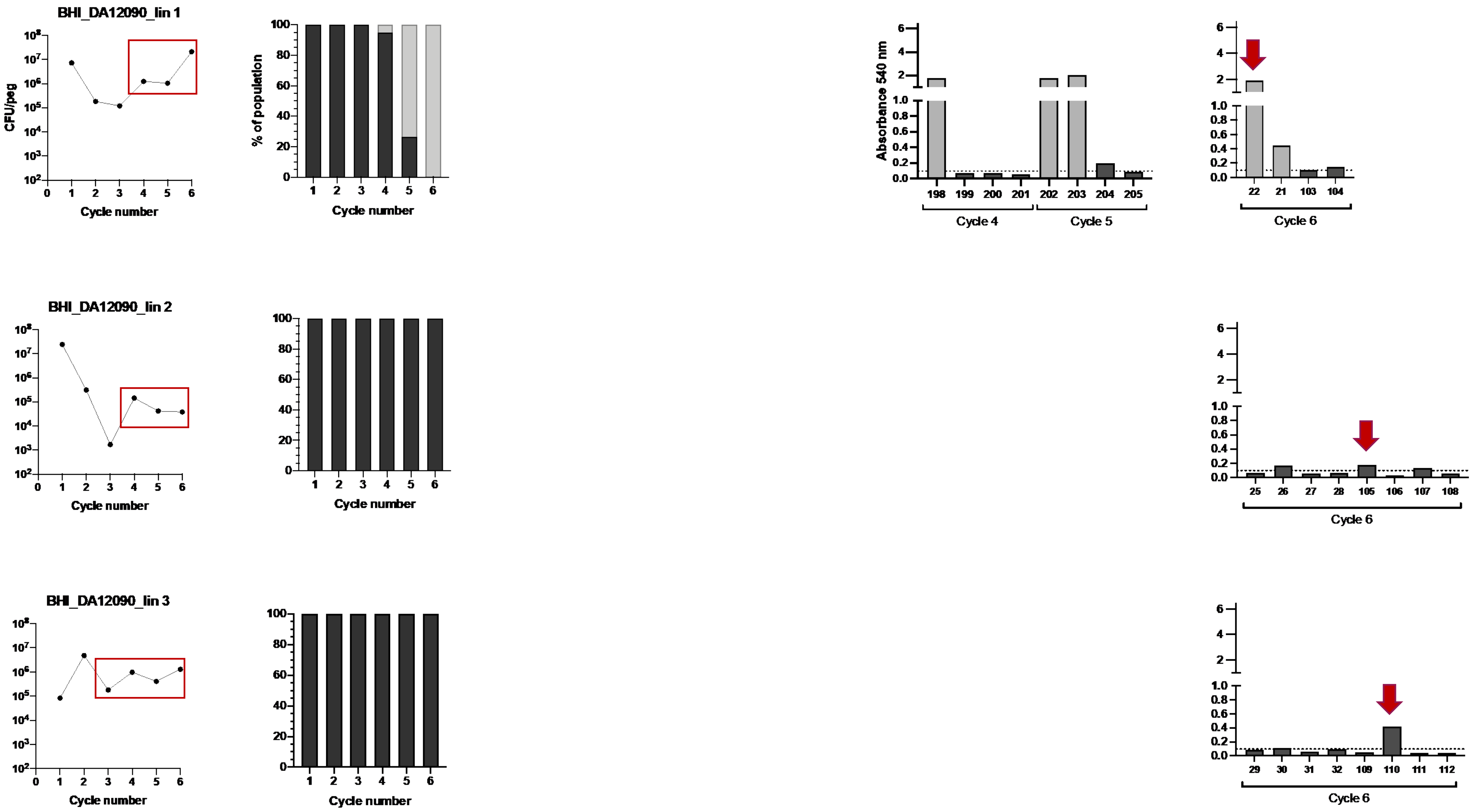
BHI_DA12090_lin 1
BHI_DA12090_lin 2
BHI_DA12090_lin 3
CFU/peg
Cycle number
% of population
Absorbance 540 nm
Cycle 4
Cycle 5
Cycle 6

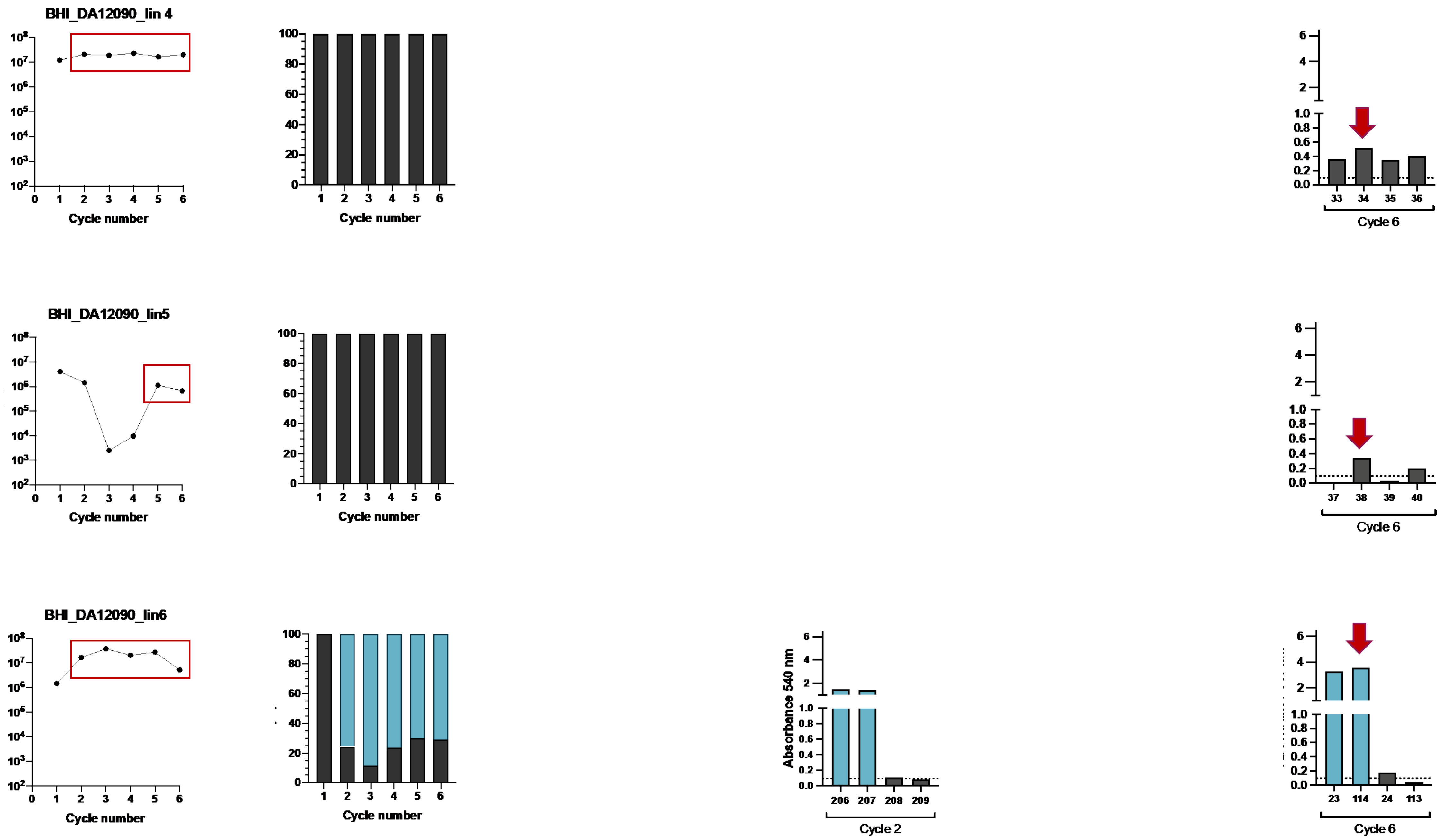
BHI_DA12090_lin 4
Cycle number
Cycle number
33
34
35
36
Cycle 6
BHI_DA12090_lin5
Cycle number
Cycle number
37
38
39
40
Cycle 6
BHI_DA12090_lin6
Cycle number
Cycle number
Absorbance 540 nm
206
207
208
209
Cycle 2
23
114
24
113
Cycle 6

### BHI_DA14734_lin1

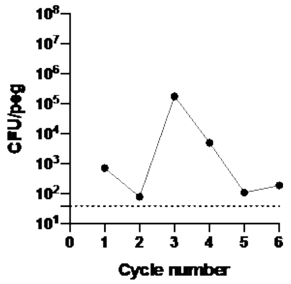

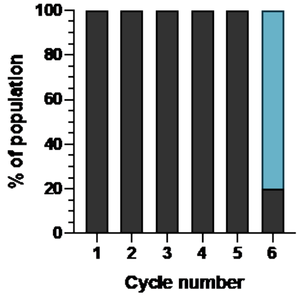

### BHI_DA14734_lin2

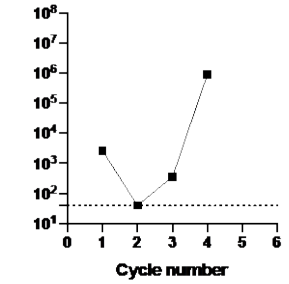

### BHI_DA14734_lin3

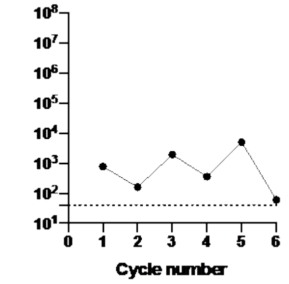

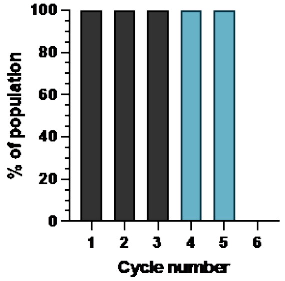

BHI_DA14734_lin 4

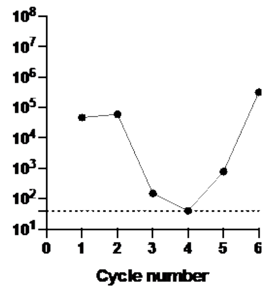

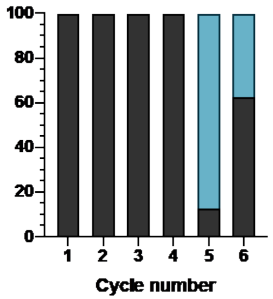

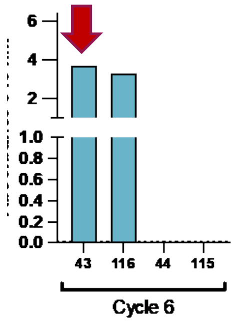

BHI_DA14734_lin 5

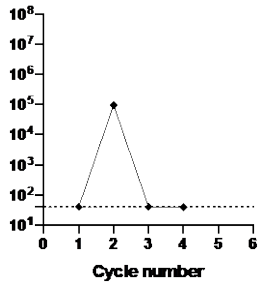

BHI_DA14734_lin 6

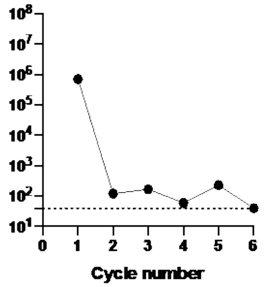

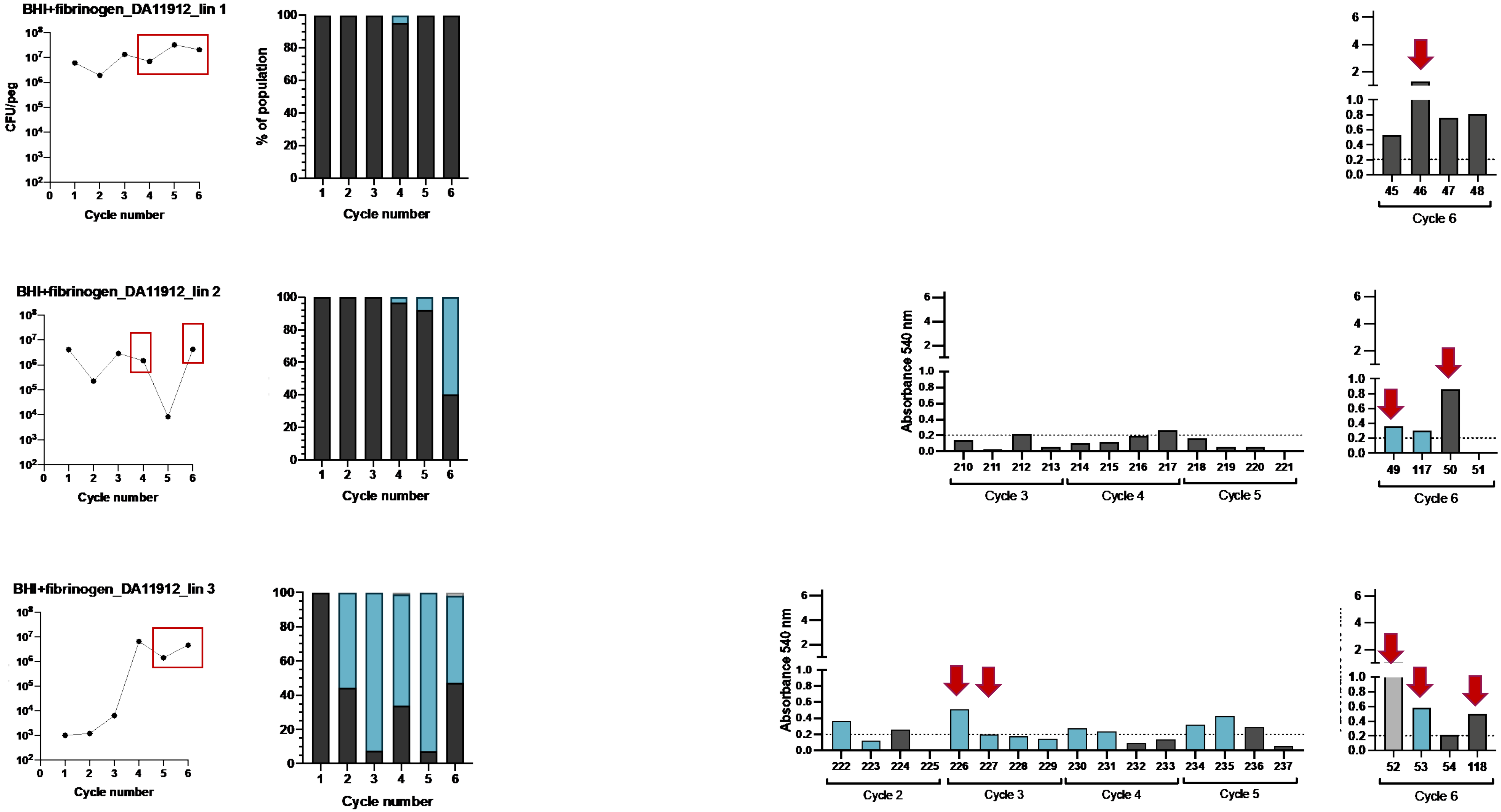
BHI+fibrinogen_DA11912_lin 1
CFU/peg
Cycle number
% of population
Cycle number
45 46 47 48
Cycle 6
BHI+fibrinogen_DA11912_lin 2
Cycle number
Cycle number
Absorbance 540 nm
210 211 212 213 214 215 216 217 218 219 220 221
Cycle 3
Cycle 4
Cycle 5
49 117 50 51
Cycle 6
BHI+fibrinogen_DA11912_lin 3
Cycle number
Cycle number
Absorbance 540 nm
222 223 224 225 226 227 228 229 230 231 232 233 234 235 236 237
Cycle 2
Cycle 3
Cycle 4
Cycle 5
52 53 54 118
Cycle 6

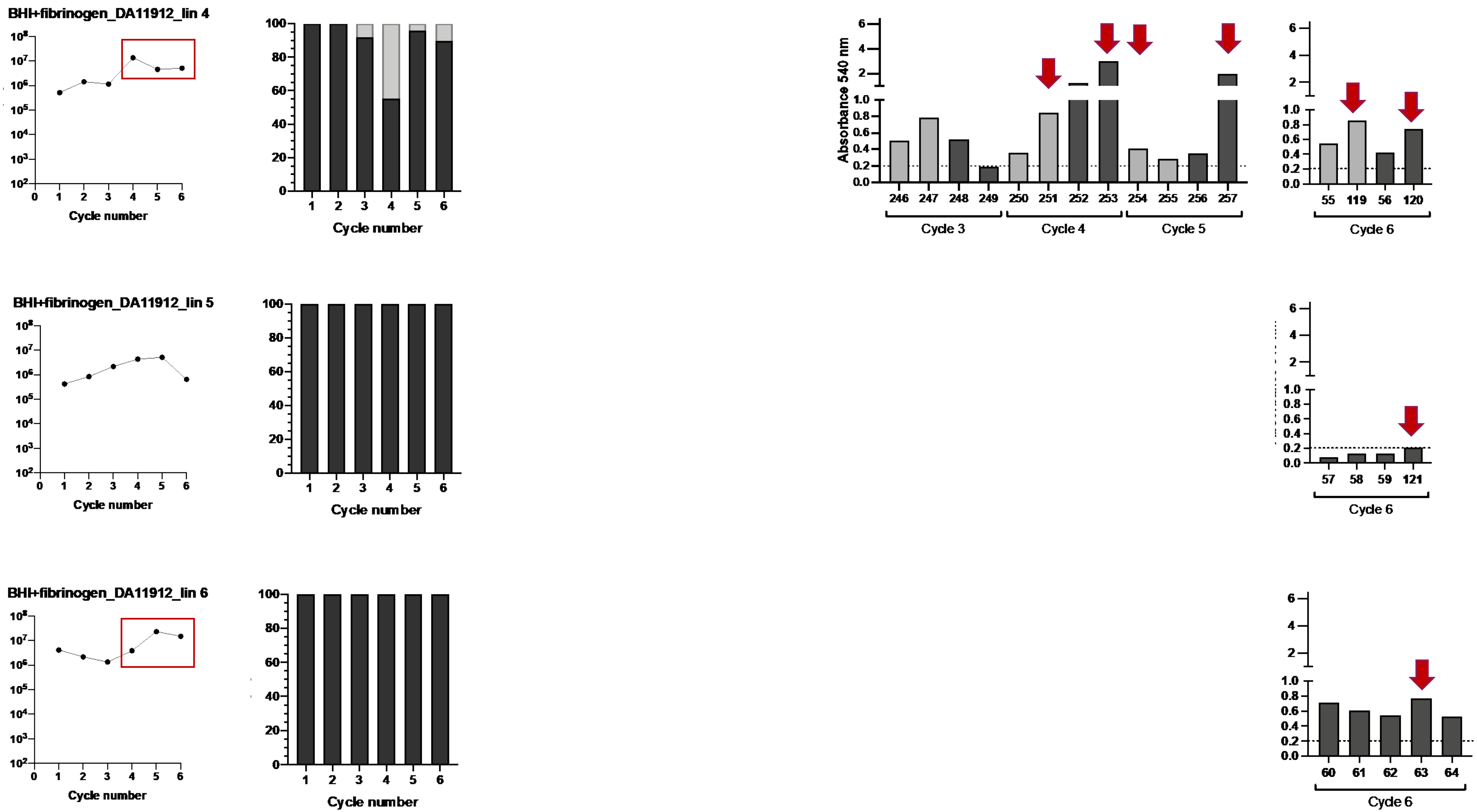
BHI+fibrinogen_DA11912_lin 4
BHI+fibrinogen_DA11912_lin 5
BHI+fibrinogen_DA11912_lin 6
Cycle number
Absorbance 540 nm
Cycle 3
Cycle 4
Cycle 5
Cycle 6

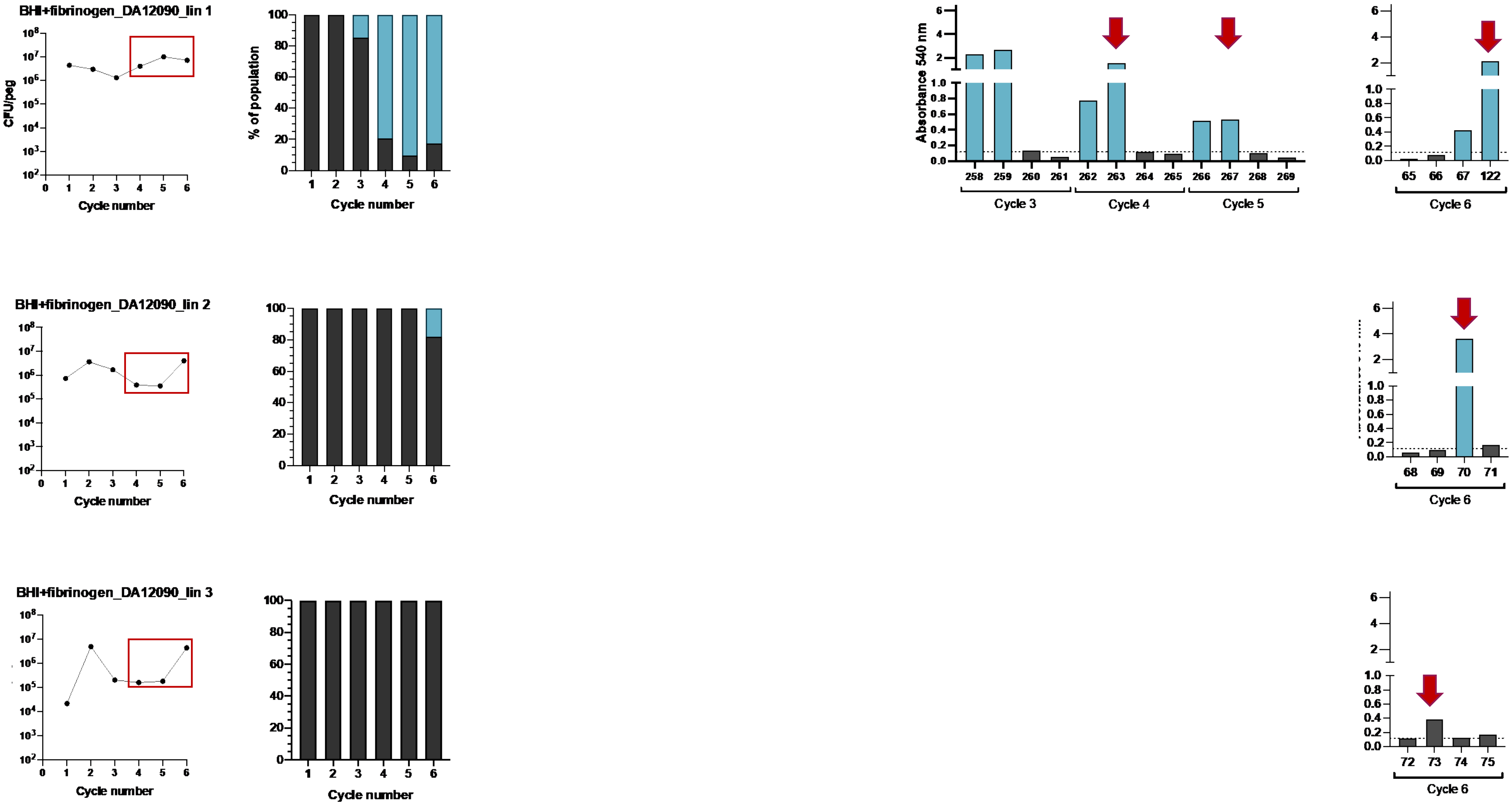
BHI+fibrinogen_DA12090_lin 1
BHI+fibrinogen_DA12090_lin 2
BHI+fibrinogen_DA12090_lin 3
CFU/peg
% of population
Cycle number
Absorbance 540 nm
Cycle 3
Cycle 4
Cycle 5
65
66
67
122
68
69
70
71
72
73
74
75
Cycle 6

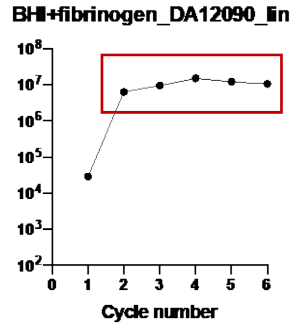
BHI+fibrinogen_DA12090_lin 4
Cycle number
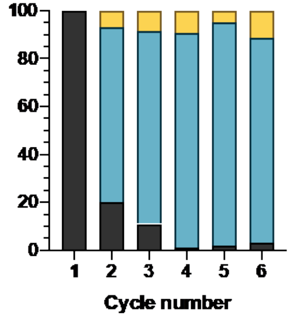
Cycle number
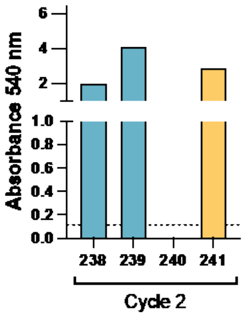
Absorbance 540 nm
238
239
240
241
Cycle 2
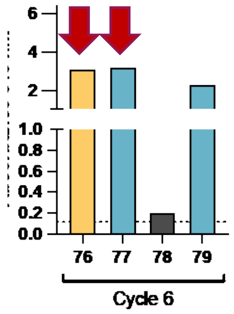
76
77
78
79
Cycle 6
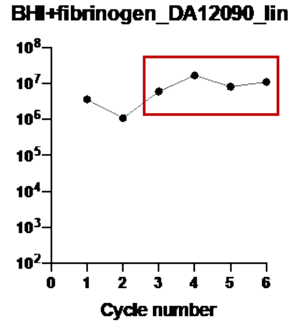
BHI+fibrinogen_DA12090_lin 5
Cycle number
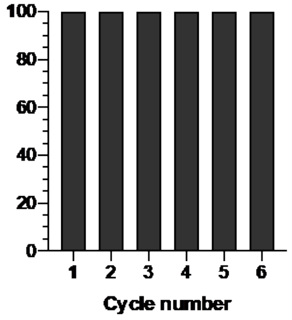
Cycle number
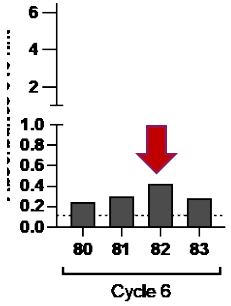
80
81
82
83
Cycle 6
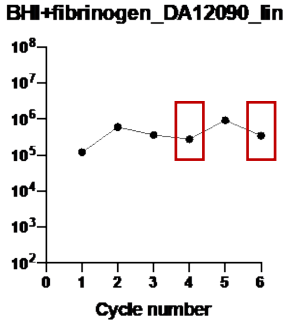
BHI+fibrinogen_DA12090_lin 6
Cycle number
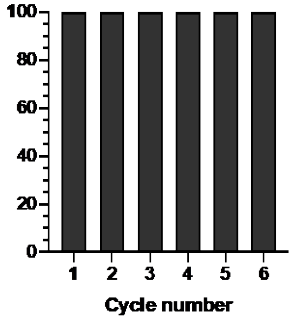
Cycle number
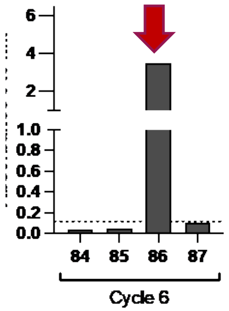
84
85
86
87
Cycle 6

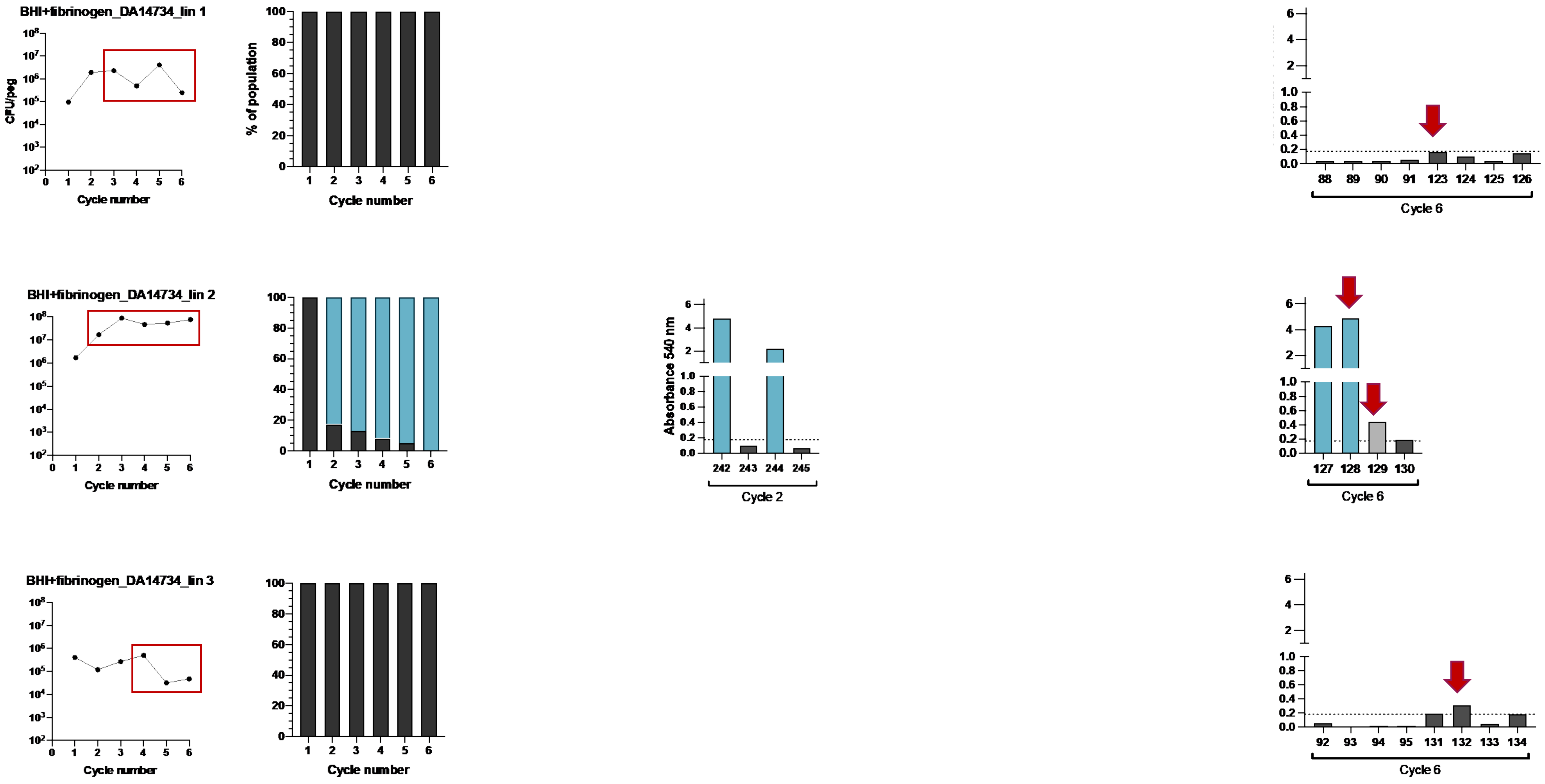

BHI+fibrinogen_DA14734_lin 1
CFU/peg
Cycle number
% of population
Cycle number
88
89
90
91
123
124
125
126
Cycle 6
BHI+fibrinogen_DA14734_lin 2
Cycle number
Cycle number
Absorbance 540 nm
242
243
244
245
Cycle 2
127
128
129
130
Cycle 6
BHI+fibrinogen_DA14734_lin 3
Cycle number
Cycle number
92
93
94
95
131
132
133
134
Cycle 6

BHI+fibrinogen_DA14734_lin 4

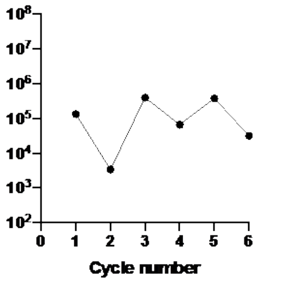

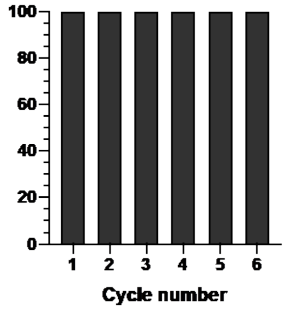

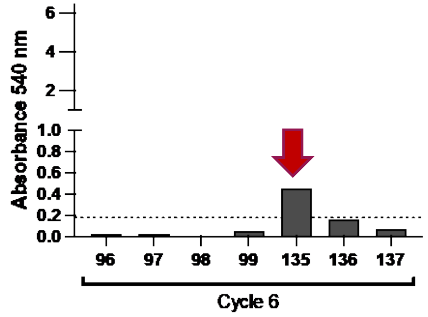

BHI+fibrinogen_DA14734_lin 5

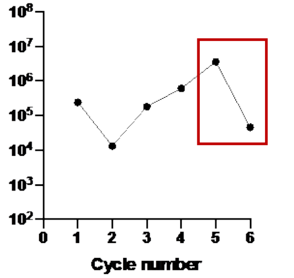

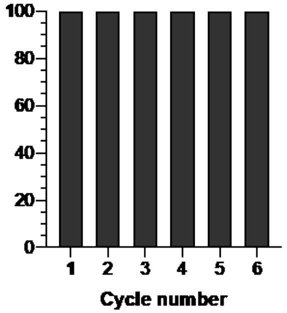

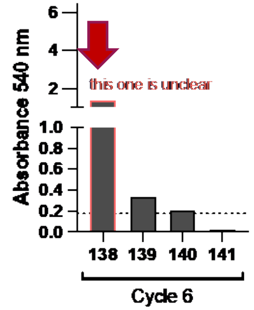

BHI+fibrinogen_DA14734_lin 6

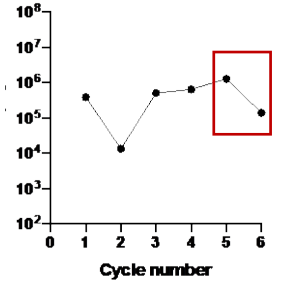

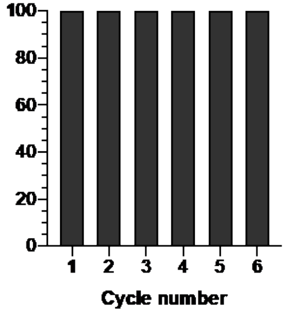

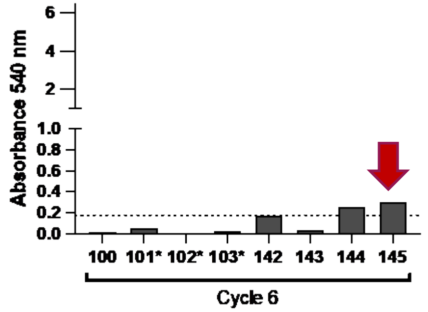
