## Supplementary Figures for "Rapid evolution of *Klebsiella pneumoniae* biofilms *in vitro* delineates adaptive changes selected during infection"

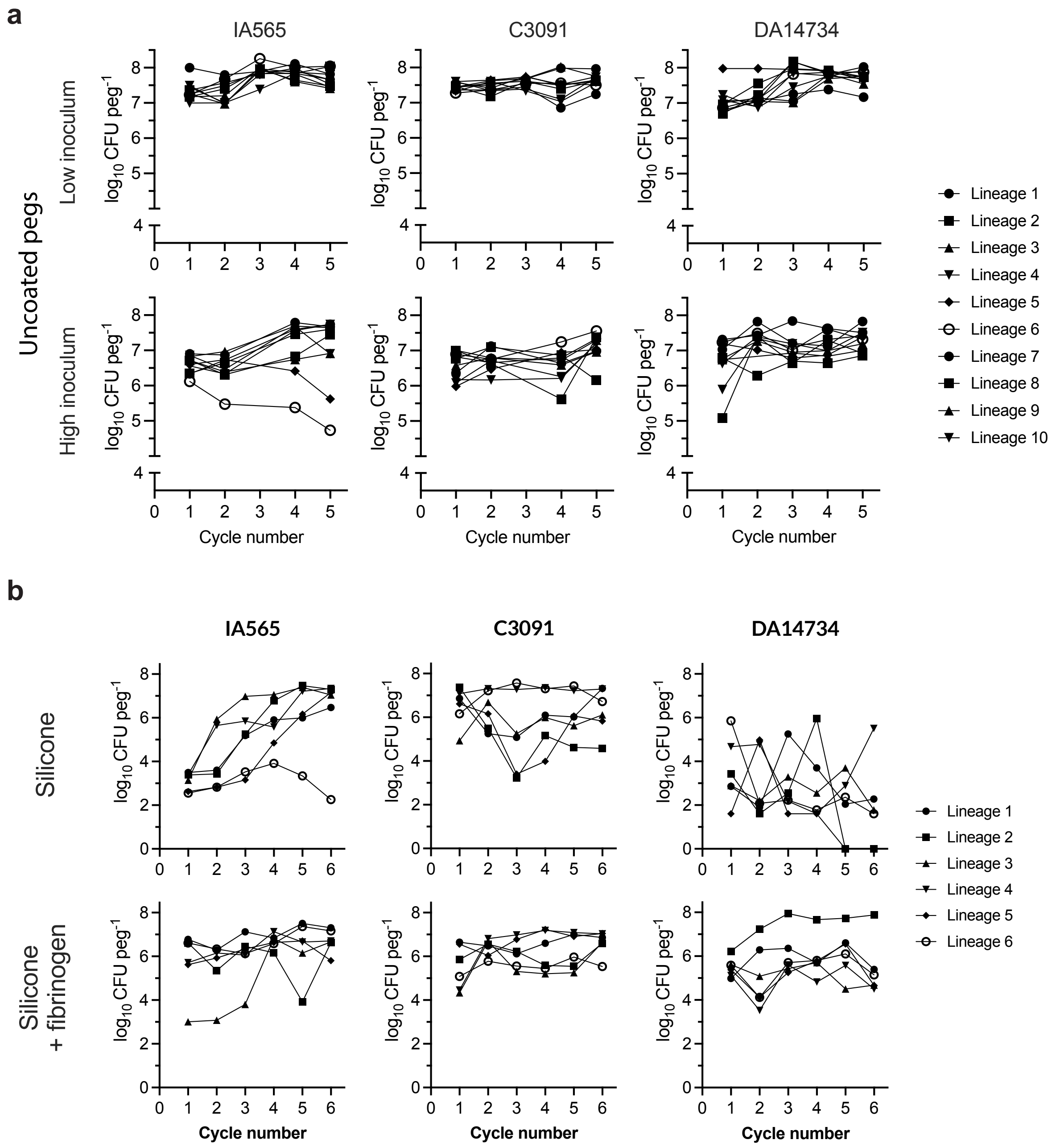


**Supplementary Fig. 1 |** Population size changes during serial passaging on **a** uncoated pegs and **b** silicone-coated pegs with (bottom) and without (top) fibrinogen during serial passaging in BHI medium.

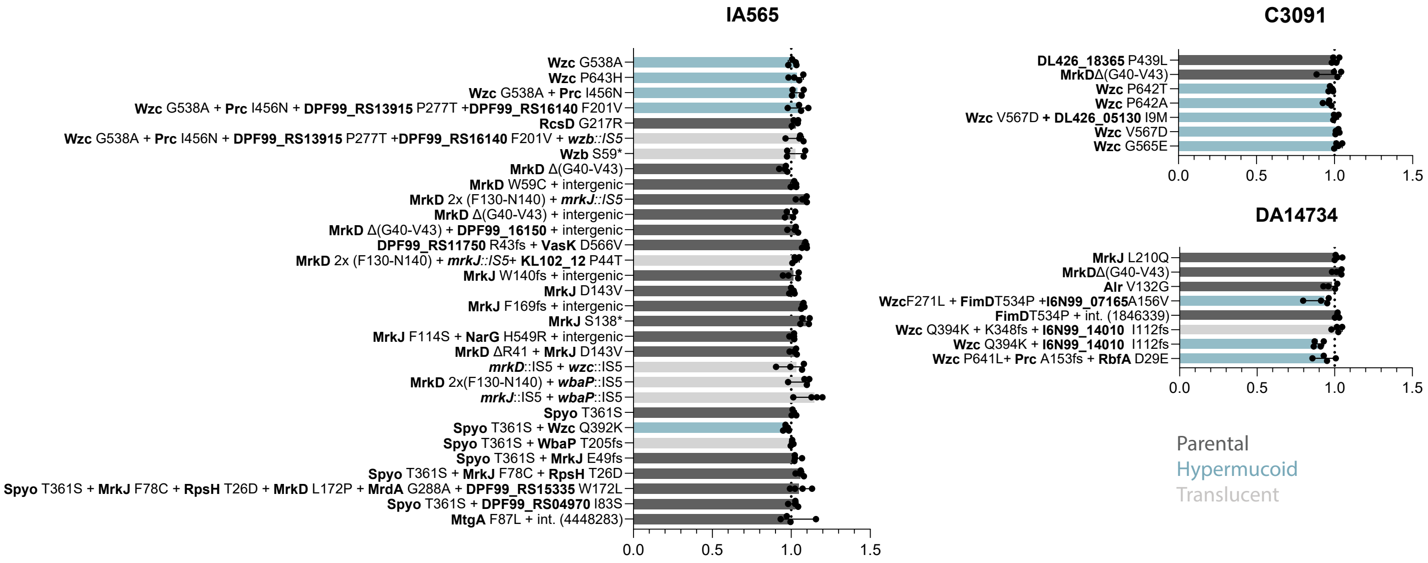


**Supplementary Fig. 2 |** Exponential growth rates in biofilm-evolved mutants during planktonic growth in BHI relative to the parental strain.

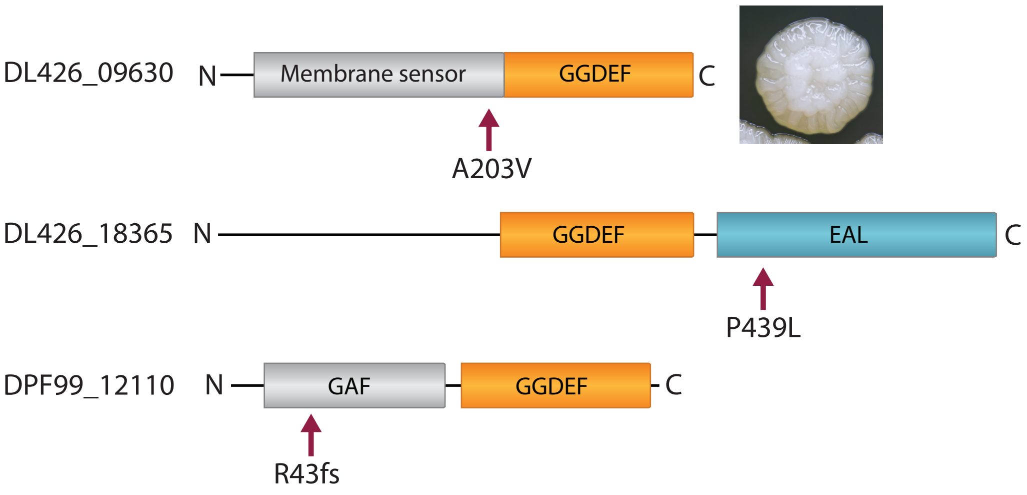


**Supplementary Fig. 3 |** Protein domain structures based on protein BLAST results in uncharacterized EAL/GGDEF proteins from IA565 (DPF99_12110) and C3091 (DL426_18365 and DL426_09630). Arrows mark the mutations. A203V in DL426_09630 leads to a unique wrinkly colony morphology.

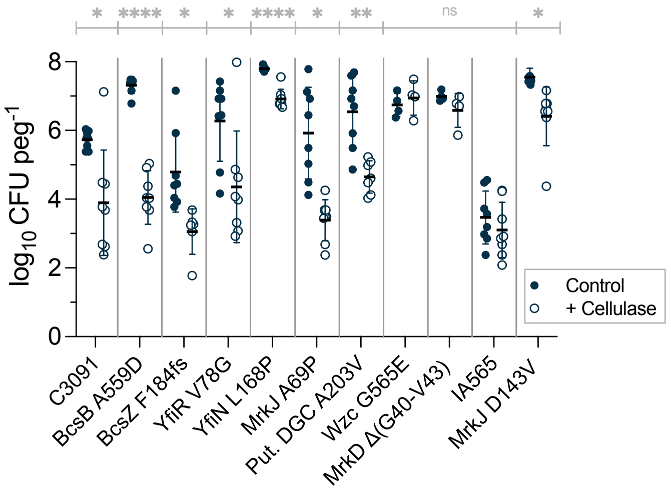


**Supplementary Fig. 4 |** Biofilm growth on silicone-coated pegs (24 h) in the presence of cellulase enzyme. Results from eight biological replicates with 95% CI are shown. Statistical significance was assessed by a paired two-tailed Student's t-test between control and treated biofilms for each strain.
